# Can Antisense lncRNA bind to the product of its sense pair?

**DOI:** 10.64898/2026.01.23.701224

**Authors:** Nair Sreejith, Singh Deepshikha, Saha Aditi, Datta Bhaskar, Majumdar Sharmistha

## Abstract

Long non-coding RNAs (lncRNAs) account for a major proportion of the transcriptional output in complex organismal genomes. Their emergence as auxiliary regulators of gene expression as well as their roles in metastasis and cancer progression has put them in the limelight. LncRNAs perform multitudes of functions and often moonlight as regulators, scaffolds and guides. Most lncRNAs are cell and tissue specific and can act as markers for diseases as well as targets for therapeutic interventions. LncRNAs are also known to make use of higher order structures such as G-quadruplexes (G4) to facilitate complex functions and interactions. THAP9-antisense1 (AS1) is a lncRNA coding gene (recently annotated by Ensembl) that codes for 12 lncRNA transcripts and has been implicated in many disease pathologies like gastric cancer, spontaneous neutrophil apoptosis, hepatocellular carcinoma, and the progression of oesophageal cancer. It is the antisense gene pair of the THAP9 gene ( a transposase derived gene) with which it shares a promoter. THAP9-AS1 has been reported to be dysregulated during stress and several cancers. However, the exact role of the lncRNA is not well understood. Bioinformatics driven strategies are used to identify putative quadruplex forming sequences (PQSs) within the lncRNA THAP9-AS1. The identified PQSs are further validated using biophysical, spectroscopic and molecular biology driven techniques. The importance of each G-tract in the formation of a particular RNA G-quadruplex (rG4) is studied via the investigation of several deletion mutants. The findings demonstrate the rG4 forming potential of the identified PQSs within THAP9-AS1.

## Introduction

For many decades, RNA has been viewed primarily as an intermediate molecule between DNA and protein, only serving as a messenger RNA (mRNA). A change in this view occurred with the discovery of a large fraction of the genome which does not encode for proteins. This discovery along with the discovery of regulatory roles of RNA, led to these RNAs being termed as non-coding RNAs (ncRNAs) (Mattick et al., 2023a).

Among the ncRNAs those >200 Nts in length were termed as long non-coding RNAs (lncRNAs) (Mattick et al., 2023b). This size threshold distinguishes them from small ncRNAs and miRNAs, siRNAs (Zhang et al., 2023) and piRNas (Haase et al., 2024). These lncRNAs are known to play a regulatory role in the cell.

The concept of functional lncRNAs predates modern genomics and can be traced back to discovery of H19 (Pachnis et al., 1984) and XIST in mammals (Brown et al., 1992), roX1 and roX2 in *Drosophila* (Meller & Rattner, 2002).

Until the discovery of their regulatory roles, these transcripts were considered to be an anomaly in the cell and termed as Junk. High throughput transcriptomics have now shed light on various lncRNAs present with cell type specific expression patterns which has lead to overturning the notion of them being Junk part of the genome and establishing them as regulators of gene expression.

lncRNAs based on their diverse genomic origins can be classified as (i) intergenic lncRNA (Cabili et al., 2011) (ii) intronic lncRNA (Rinn & Chang, 2012) (iii) bidirectional lncRNA sharing promoter with coding genes (Albrecht & Ørom, 2016) (iv)antisense lncRNA (Pelechano & Steinmetz, 2013) (v) enhancer RNAs (Woolard & Chorley, 2018) and (vi) overlapping lncRNAs (Katayama et al., 2005).

lncRNAs are transcribed by RNA polymerase II and undergo 5’ capping and polyadenylation (Bhat et al., 2016). lncRNAs exhibit lower expression levels and high tissue specificity (Grammatikakis & Lal, 2021). lncRNAs undergo rapid and volatile evolution and show limited sequence conservation (Qu & Adelson, 2012). Often functional conservation is dependent on the structure rather than the sequence, emphasizing on the importance of secondary and tertiary structures.

lncRNA functions as guides, scaffolds, decoys, signals and enhancers (Fang & Fullwood, 2016). lncRNAs regulate chromatin organization, transcription, post-transcriptional processing and nuclear architecture via these modes (Fernandes et al., 2019).

One of the examples of lncRNA’s role in chromatin remodeling and epigenetic regulation is that of X-chromosome inactivation by XIST lncRNA. The XIST lncRNA inactivates the expression of X-chromosome via recruiting the PRC2 complex and promoting H3K27me3 deposition on the chromosome (Zhao et al., 2008).

Similarly, HOTAIR functions as a modular scaffold by coordinating PRC2 and LS1 to silence expression of the HOX gene (Wu et al., 2015).

lncRNAs are known to modulate the enhancer-promoter looping, recruit transcription factors and regulate RNA polymerase II activity(T. K. Kim et al., 2010). lncRNAs are also known to influence alternative splicing as well as act as competing endogenous RNAs as well as sequester RNA-binding proteins(Gerstberger et al., 2014; Hou et al., 2023; Salmena et al., 2011).

### RNA G-quadruplexes (rG4s) in lncRNA

G-quadruplexes are non-canonical nucleic acid structures formed via stacking of G-tetrads and stabilized by monovalent cations (Vorlíčková et al., 2007). rG4s, were first described in 1991 ( Kim et al., 1991) and are known to be more compact and thermodynamically stable than their DNA counterparts. They are known to predominantly adopt parallel conformations (Zaccaria & Fonseca Guerra, 2018).

rG4 motifs are widespread in mRNAs and lncRNAs and can be detected via spectroscopic techniques, chemical probing and high throughput approaches (Santos et al., 2021). Many lncRNAs are rich in G-rich regions and this helps in the formation of rG4 motifs. These rG4 motifs help modulate RNA stability, localization and protein interaction (Statello et al., 2021). Terra is a classic example of lncRNA whose rG4 motif recruits TRF2 and regulates telomere function (Mei et al., 2021). rG4 motif present in NEAT1 mediates the recruitment of NONO and SFPQ, thereby driving assembly of paraspeckles (Simko et al., 2020).

### Mechanisms of lncRNA-protein interaction

lncRNAs are known to interact with proteins via sequence motifs, secondary and tertiary structures and multivalent cooperative binding (Cléry et al., 2013; Fang et al., 2024). Structural motifs such as stem loops, bulges, triple helices, G-quadruplexes are known to serve as protein docking stages (Hermann & Patel, 2000; Obuse & Hirose, 2023).

Helicases (e.g., DHX36, WRN), RBPs, Transcription factors and chromatin modifiers are known to bind to rG4s (Booy et al., 2012; Meier-Stephenson, 2022). In the case of lncRNAs like HOTAIR and HOXD-AS1, EZH2 binds preferentially to rG4 present on the lncRNA and has a tumor suppressive effect (Hao et al., 2021).

rG4 mediated interactions are also known to induce allosteric regulation of proteins. DHX36-GSEC and hnRNP F-CD44 systems are the known examples where such regulation is seen.

### THAP9-AS1

THAP9-AS1 is a newly annotated lncRNA transcribed from the antisense gene pair of the THAP9 gene, with which it shares a bidirectional overlapping promoter. Pan-cancer analysis reveals THAP9-AS1 is dysregulated across various cancers and is often correlated with poor prognosis(Rashmi & Majumdar, 2022).

THAP9-AS1 participates in oncogenic feedback in pancreatic ductal adenocarcinoma (PDAC) and Oesophageal squamous cell carcinoma (ESCC) via miRNA sponging and YAP/SOX4 signalling(Cheng et al., 2021; Li et al., 2020). THAP9-AS1 is also implicated in HCC (Su et al., 2022), gastric cancer (Jia et al., 2019), and pediatric septic shock (Wu et al., 2020).

The protein interactome and structural information of the lncRNA however remains largely unexplored.The presence of a shared promoter between the THAP9 and THAP9-AS1 hints at a regulatory role that could be at play. Given the importance of rG4 in mediating interaction between various interactomes in the cell, the study explores putative G-quadruplex forming sequence (PQS) within THAP9-AS1 and its possible interaction with THAP9 protein.

## 2. Results

### 2.1 Prediction of PQS in lncRNA THAP9-AS1

THAP9-AS1 lncRNA has recently been implicated in various cancers (Su et al., 2022). However, not much is known about the lncRNA’s mechanism of action, biological role or mode of interaction. It is known that lncRNAs interact with other cellular partners via secondary structures like G4 motifs. Thus, we investigated the propensity of THAP-AS1 lncRNA to form rG4 structures.

ENSEMBL lists three (out of 12) transcript variants (Table 1) with verified Refseq status for lncRNA THAP9-AS1 (ENSG00000251022). The three transcript variants have varying lengths [transcript variant 1 (NR_034075.1): 2,318 base pairs, transcript variant 2 (NR_034076.1): 2,293 base pairs, transcript variant 3 (NR_034077.1): 2,219 base pairs https://www.ncbi.nlm.nih.gov/datasets/gene/id/100499177/products/].

**Table 1.** Transcript variants of THAP9-AS1 and their NCBI accession ID.

| **Transcript** | **Accession ID** | **Length (bp)** |
| --- | --- | --- |
| **THAP9-AS1 variant 1** | **NR_034075.1** | **2,318** |
| **THAP9-AS1 variant 2** | **NR_034076.1** | **2,293** |
| **THAP9-AS1 variant 3** | **NR_034077.1** | **2,219** |

The quadruplex forming ability of each THAP9-AS1 transcript variant (Table 1) was predicted using different G4 finder tools namely imGQfinder (Varizhuk et al., 2014), G4 hunter (Brázda et al., 2019)and QGRS mapper (Kikin et al., 2006). imGQfinder did not identify any PQS in THAP9-AS1 transcript variants (1-3). All three transcript variants were analyzed using G4 hunter with a sliding window ranging from 20-40. However, no G4s were predicted.

Interestingly, QGRS mapper predicted more than one PQS (Putative Quadruplex forming Sequence) in each transcript variant (Table 2). These results were selected over that of imGQ and G4 Hunter as QGRS mapper is a widely used tool that allows users to define loop length, number of tetrads and motif length (Frees et al., 2014).

**Table 2.**
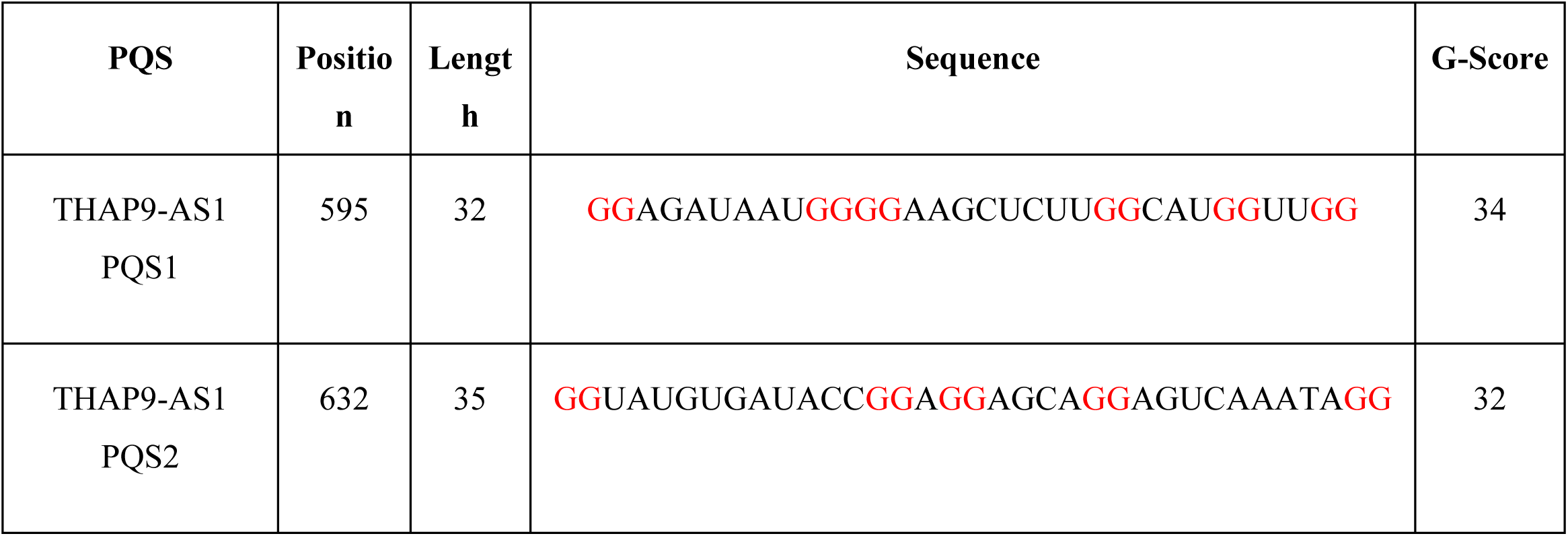
Putative Quadruplex forming sequences (PQS, G tracts in red font) obtained from QGRS mapper, present across all 3 transcript variants of lncRNA THAP9-AS1 (representative image from results of PQS1 and PQS2 in THAP9-AS1 Transcript Variant 3)

Two PQSs which were predicted in all three transcript variants (Table 2) were selected for further analysis. Figure S5. shows the relative position of the PQSs identified within the THAP9-AS1 gene. Table 5 also shows the sequence position in THAP9-AS1 transcript variant 3 for PQS1 (595-629 nt) and PQS2 (632-664 nt).

### 2.2 *In vitro* characterization of PQS identified in lncRNA THAP9-AS1

#### 2.2.1 Native Gel visualization of PQS1 and PQS2 and deletion mutants

The PQS1 and PQS2 identified in the lncRNA THAP9-AS1 have five G tracts. PQS1 is made up of three 2G tracts and one 4G tract, whereas, PQS2 is composed entirely of 2G tracts Fig. 1). To determine the importance of individual G tracts (red font, Table S4) in maintaining the structure of each PQS, a series of deletion mutants were created by sequentially deleting each G-tract (Fig. 1).

**Figure 1.**
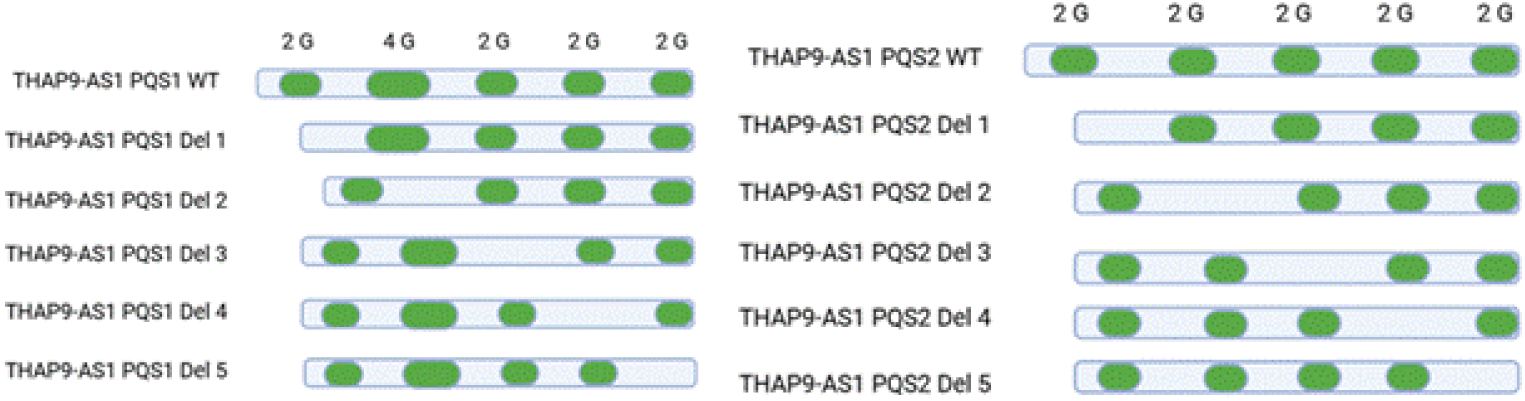
Schematic of PQS1 and PQS2 Wild type and deletion mutants

PQS1 and PQS2 and their respective deletion mutants were synthesized via *in vitro* transcription using the corresponding antisense DNA sequence as template. A native gel (Fig. 2) was run to check for the presence of G4 formation that was identified by ThT staining.

**Figure 2.**
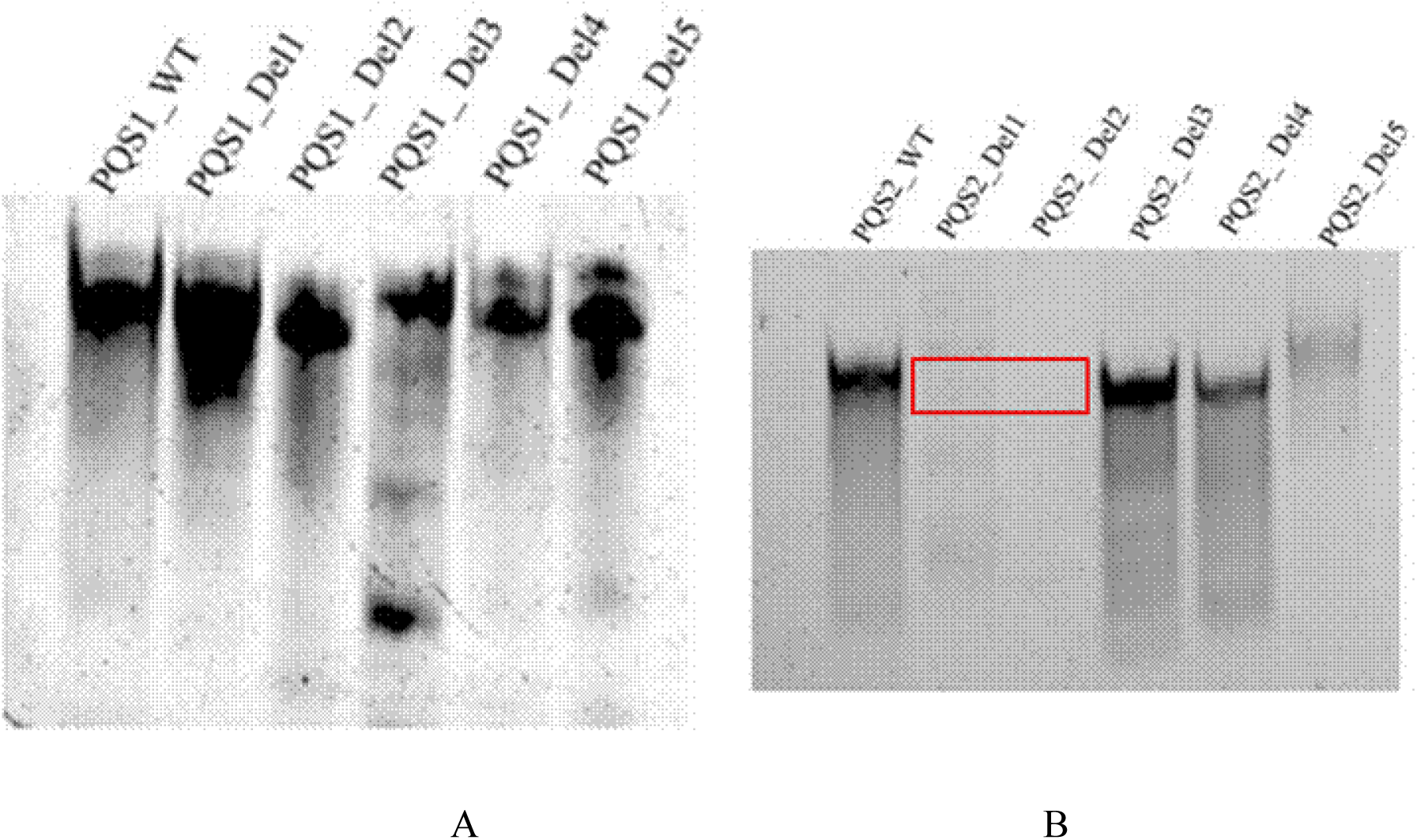
Native gel image of (A) PQS1 and deletion mutants, (B) PQS2 and deletion mutants stained with ThT to check for presence of G4

The ThT stained-native gel image of PQS1 and its deletion mutants (Fig. 2 (A)) illustrates G4 formation in the wild type sequence (PQS1_WT) as well as the deletion mutants. On the other hand, PQS2 Deletion mutants 1 and 2 (Fig. 2 (B)) undergo loss of inherent G4 formation. The G4 structure is only retained in the wild type sequence (PQS2_WT) as well as deletion mutants 3, 4 and 5 (PQS2_Del3, PQS2_Del4, PQS2_Del5).

The loss of G4 structure in PQS2_Del1 and PQS2_Del2 demonstrates the importance of specific G tracts in contributing to the stability of the G4 structure. The deletion of individual G tracts will lead to a change in the loop length between successive G tracts that may contribute to the complete loss of rG4 structure in certain mutants (marked by red box in Fig. 2) but not others. For instance, in PQS1 WT and its deletion mutants, there is no significant change in rG4 forming propensity. This suggests that even in the absence of specific G tracts, the other G tracts in PQS1 compensate and lead to formation of rG4 structure.

#### 2.2.2 CD Spectroscopy

The synthesised PQSs were then analysed by Circular Dichroism, which has been used to characterize and classify the orientation of strands in G4s (del Villar-Guerra et al., 2018).

CD spectra analysis of PQS1 and PQS2 (Fig. 3) demonstrate that they both adopt a parallel G4 structure which is an attribute of most RNA G4s, with characteristic CD maxima at 265 nm and minima at 240 nm (del Villar-Guerra et al., 2018). The CD spectra of both the PQSs were measured in the presence and absence of additional monovalent cations K^+^ and Li^+^. The parallel topology of the G4 is perceivable both in the presence and absence of the monovalent cations.

**Figure 3.**
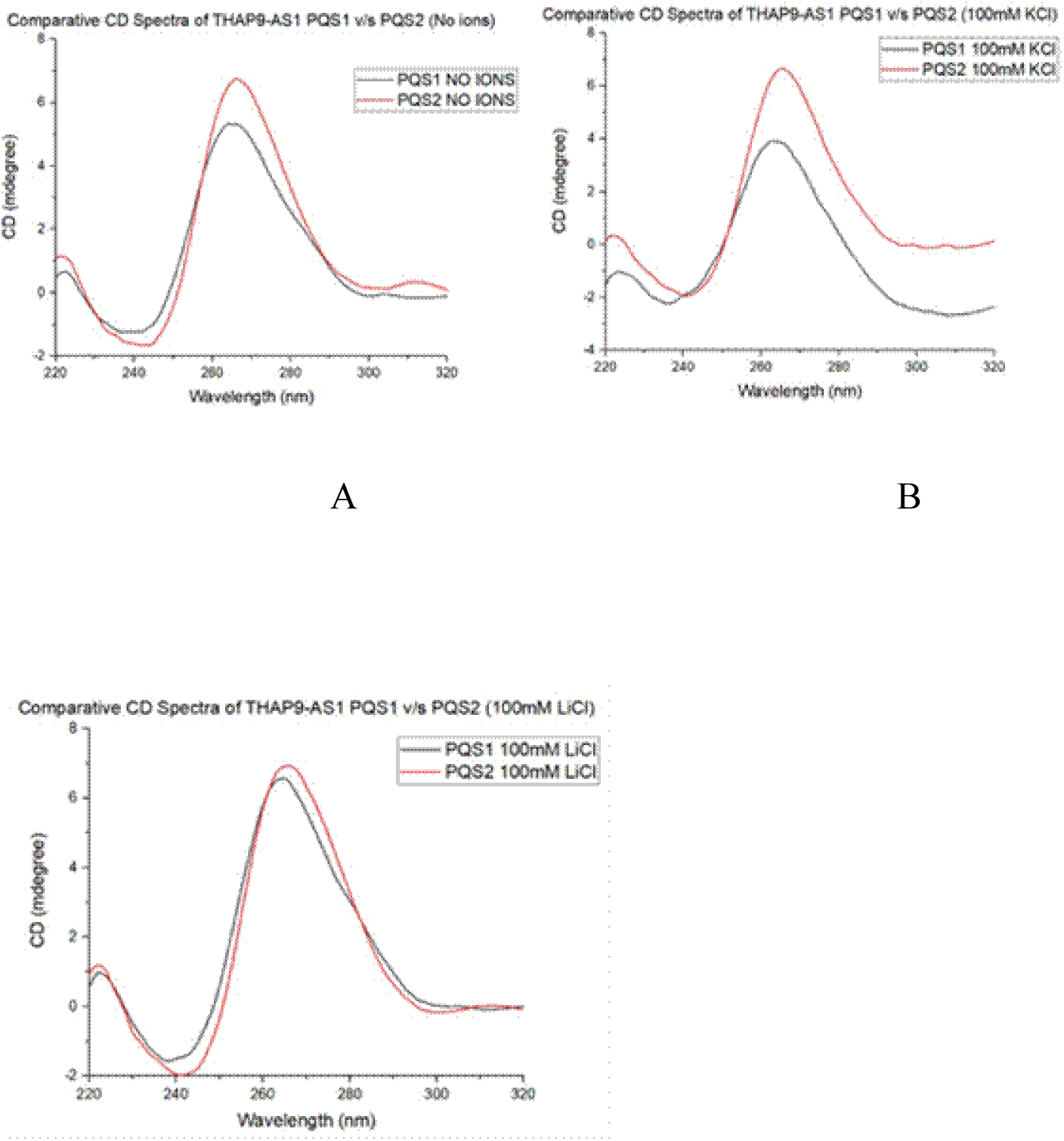
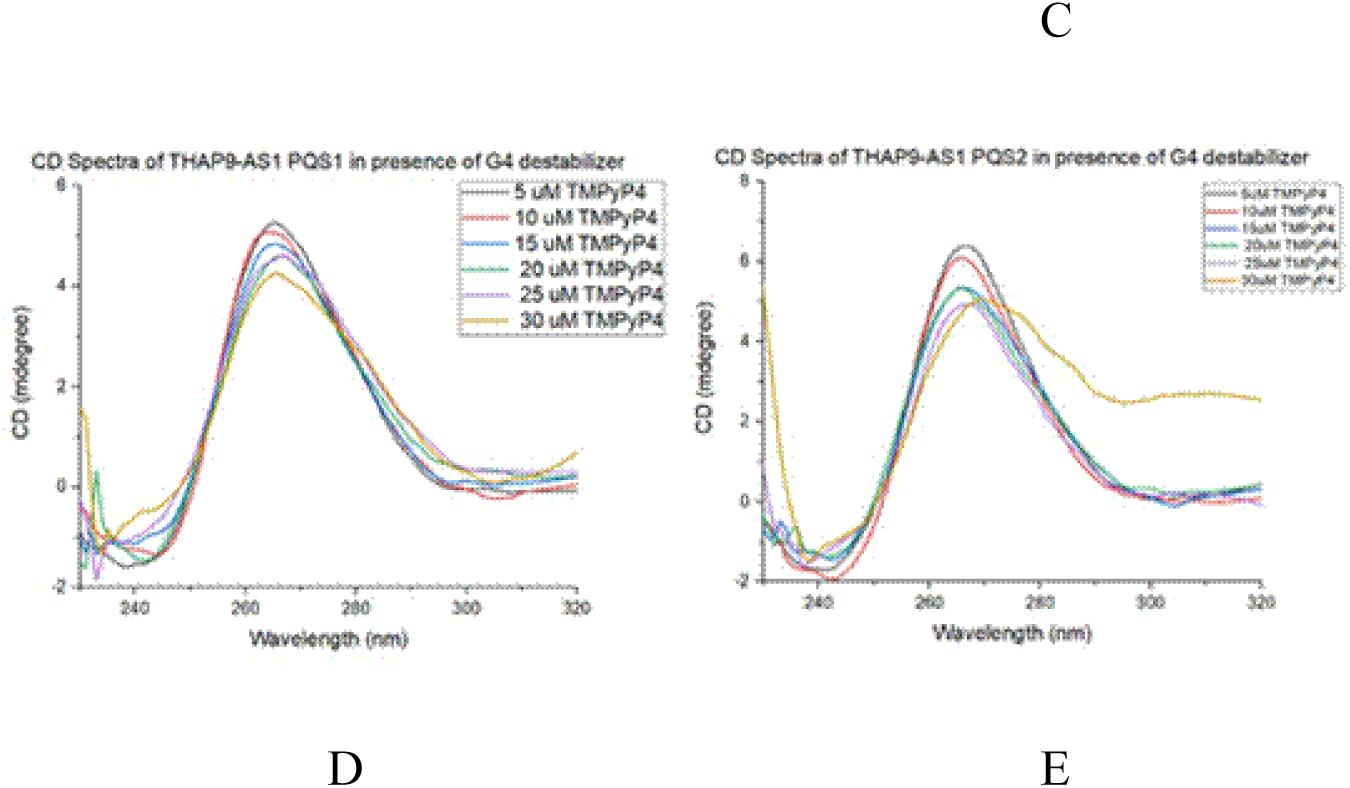
CD spectra of lncRNA THAP9-AS1 PQS1 and PQS2 (A) in absence of monovalent cations, (B) in the presence of K^+^ (100 mM), (C) in the presence of Li^+^ (100 mM), (D) comparative spectra of PQS1 in presence of increasing concentration of TMPyP4 (30 µM), (E) comparative spectra of PQS2 in presence of increasing concentration of TMPyP4 (30 µM),

It is observed that PQS2 (red) is more stable (higher maxima at 265 nm) than PQS1 (black) both in the absence (Fig. 3 A) as well as in the presence (Fig. 3 B, C) of monovalent cations. Since the CD spectra remains unchanged, it is inferred that monovalent cations do not have a pronounced effect on the G4 topology.

The stability of the G4 formed was then tested in the presence of increasing concentrations of TmPyP4 ligand, which is a cationic porphyrin that is a known G4 destabilizer. The measured CD spectra (Fig. 3 (D) and (E)) showed slight disruption (peak shouldering at 300 nm) of the parallel G4 topology at the highest concentration of TmPyP4 (30 µM). At all concentrations lower than 30 µM there appears to be no distinct effect of the ligand on the G4 structure formed.

#### 2.2.3 CD Spectroscopy of Deletion mutants of THAP9-AS1

THAP9-AS1 PQS1 comprises four 2G tracts and one 4G tract whereas PQS2 consists entirely of five 2G tracts. Deletion mutants of the respective PQSs were synthesized with successive deletion of individual G tracts.

Initial native gel analysis (Fig. 2) of PQS1 and PQS2 and their deletion mutants illustrate that for PQS1, none of the G tracts are crucial for its G4 formation potential whereas for PQS2, the deletion of the first and second G tracts leads to complete disruption and loss of G4 structure.

CD spectroscopic analysis, in the presence and absence of monovalent cations, was performed for the deletion mutants which display G4 forming potential even in the absence of one G tract. The stability of a particular mutant was correlated to higher maxima (in mdegree) at 265 nm. PQS1 deletion 1 (Fig. 4 (A)) forms a stable G4 both in the absence of ions (black) as well as in the presence of 100 mM Li^+^ ions (blue). However, its CD spectra illustrates lower stability in the presence of 100 mM K^+^ ions (red), wherein the maxima at 265 nm is relatively lower compared to the other 2 conditions. PQS1 deletion 2 (Fig. 4 (B)) shows stable G4 formation in the absence of ions (black) and shows a reverse trend with respect to the monovalent ions wherein it is more stable in the presence of K^+^ (red) as compared to Li^+^ (blue). PQS1 deletion 3 (Fig. 4 (C) shows less stable G4 formation in the absence of monovalent ions (black). In presence of K^+^ ions (red) it forms a more stable G4 compared to Li^+^ ions (blue). Both PQS1 deletion 4 (Fig.4 (D) and PQS1 deletion 5 (Fig.4 (E)) shows stable G4 formation in the absence of monovalent ions (black).

**Figure 4.**
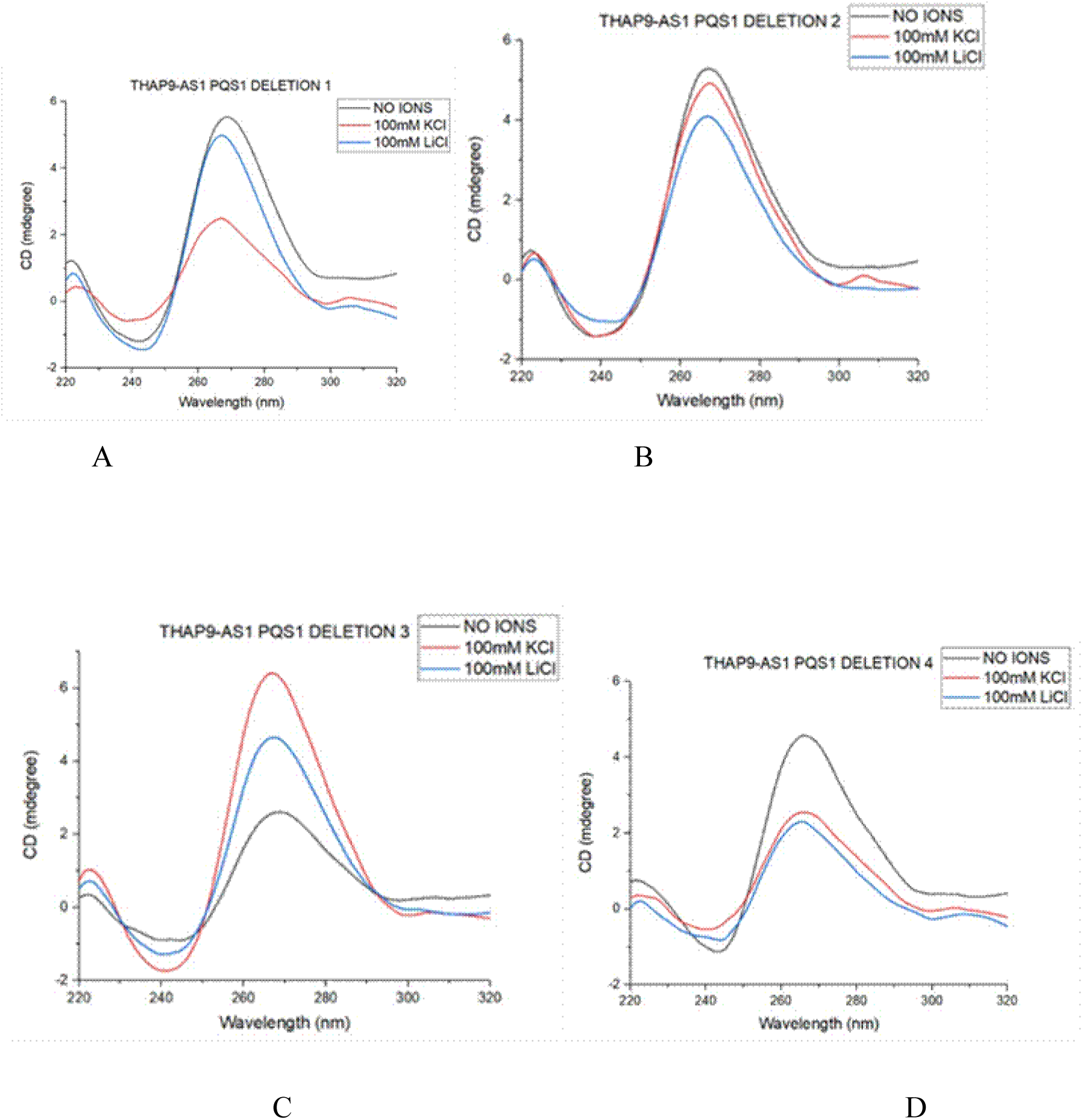
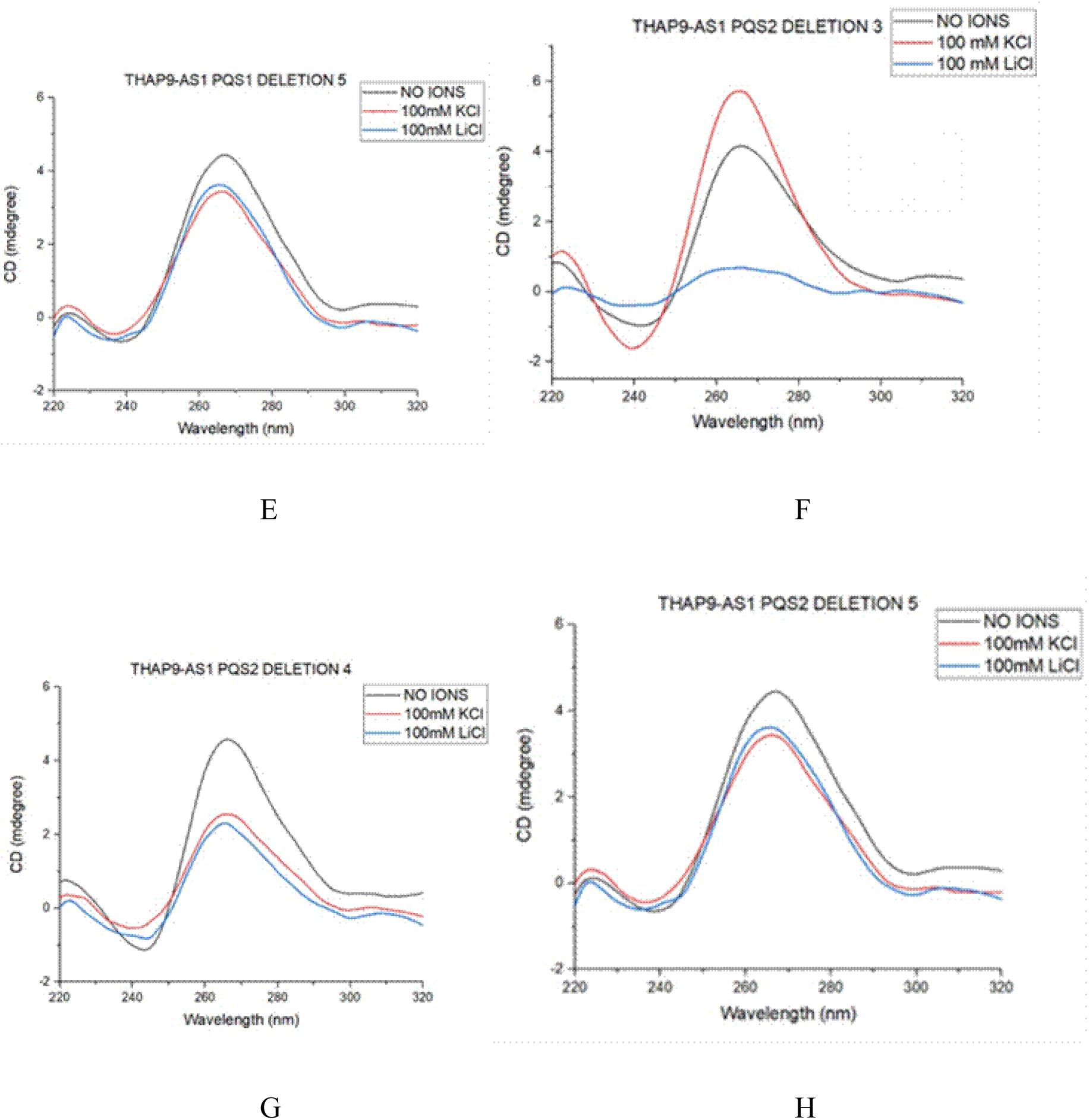
CD Spectra of Deletion mutants of PQS1 and PQS2 in the presence and absence of monovalent cations K^+^ (100 mM) and Li^+^ (100 mM) (A) PQS1 Deletion 1 (B) PQS1 Deletion 2, (C) PQS1 Deletion 3 (D) PQS1 Deletion 4 (E) PQS1 Deletion 5 (F) PQS2 Deletion 3 (G) PQS2 Deletion 4 (H) PQS2 Deletion 5. Samples were prepared in a G4-folding buffer as described in the methods section.

PQS2 deletion 3 (Fig. 4 (F) shows parallel G4 formation in the absence of monovalent ions (black) and in the presence of 100mM K^+^ ions (red). However, there is a disruption in G4 formation in the presence of 100mM Li^+^ ions (blue). In case of PQS2 deletion 4 (Fig. 4 (G)), the G4 structure formed is relatively more stable in the absence of ions (black) as compared to the presence of 100mM K^+^ (red) and Li^+^(blue). For PQS2 deletion 5 (Fig. 4 (H)), the G4 formed both in the presence and absence of monovalent ions, is of the same stability.

Thus, the native gel and CD spectra analysis of PQS1 and PQS2 demonstrate that they both adopt stable parallel rG4 structures. Such secondary structures are characteristic of most RNA G quadruplexes. The observed rG4s were often not disrupted by the presence or absence of monovalent ions as well as low concentrations of a G4 destabiliser like TmPyP4. Moreover, deletion of individual G tracts (except the first and second 2G tracts of PQS2) or competitor oligonucleotides against each G-tract (Fig. S1) mostly did not affect the stability of the rG4 structures.

#### 2.2.4 ThT Fluorescence assay

It is well reported that RNA G4 structures interact with the water soluble fluorogenic dye Thioflavin T (ThT) (Xu et al., 2016) which then undergoes an increase in fluorescence emission at 490 nm. All the PQS sequences (PQS1, PQS2 and deletion mutants) of THAP9-AS1 exhibit a characteristic ThT emission maxima at 490 nm. ThT fluorescence was enhanced 100-fold for PQS1 as compared to PQS2 (Fig. 4, 5) suggesting significant stability of the PQS1 with respect to PQS2.

Fig. 4 (A) illustrates that wild type PQS1 (red) is significantly stabler (two-fold higher fluorescence intensity, Fig. 5 (A)) than its deletion mutants both in the absence (Fig. 4 (A)) or presence of monovalent ions like K^+^ (Fig. 4 (B)) and Li^+^ (Fig. 4 (C)). Though all the PQS1 deletion mutants form stable G4s in the absence of ions (Figure 5 (A)), their G4 forming capabilities were put to test in the presence of monovalent cations. Moreover, the presence of K^+^ as compared to Li^+^, caused greater enhancement of ThT fluorescence, namely 60-fold for deletion mutant 1, 20-fold for deletion mutant 2 and 40-fold in deletion mutants 3, 4 and 5, with respect experiments done in the presence of Li^+^. This shows that K^+^ has a stabilizing effect on PQS1 deletion mutants whereas Li^+^ has an unfavourable effect.

**Figure 4.**
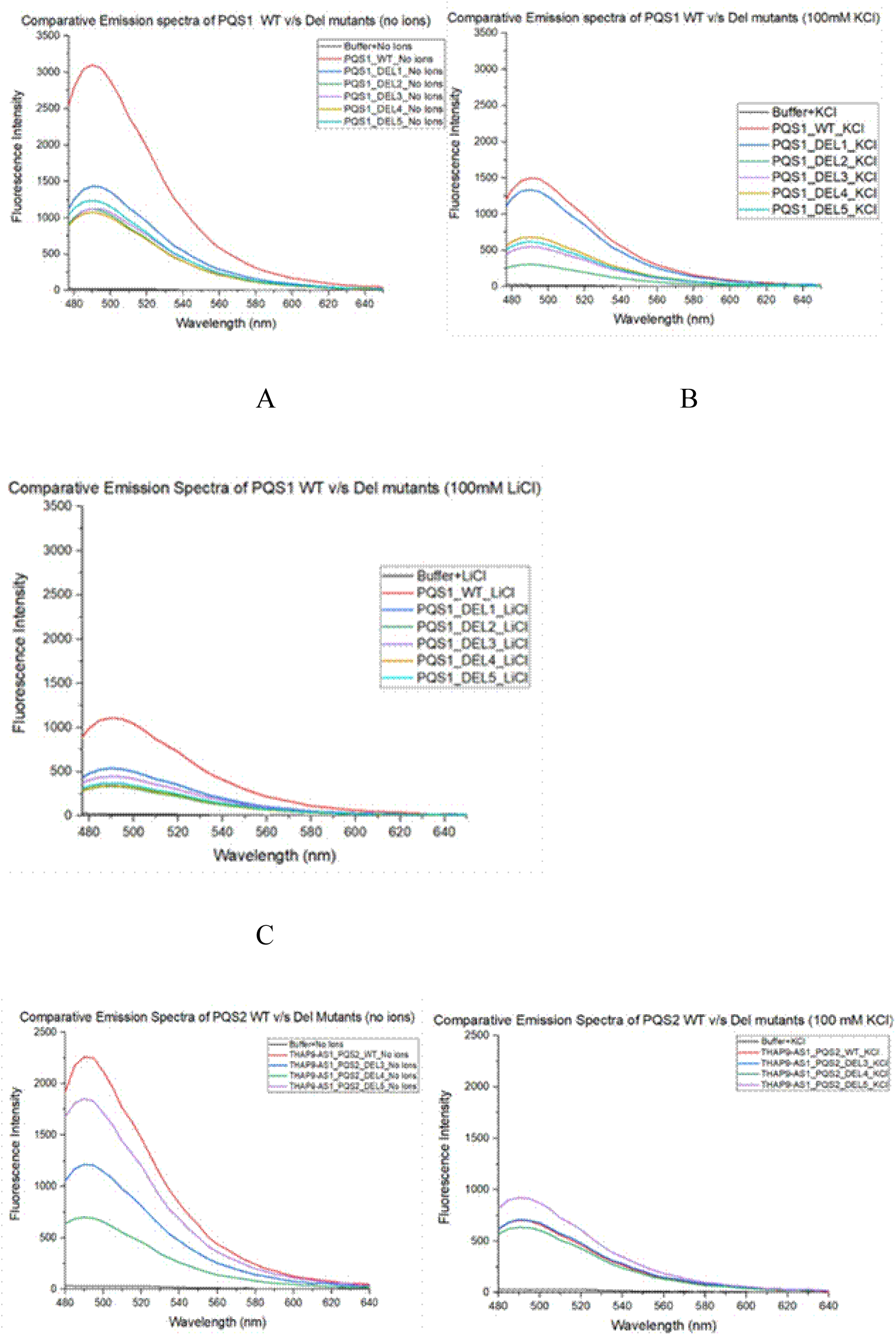
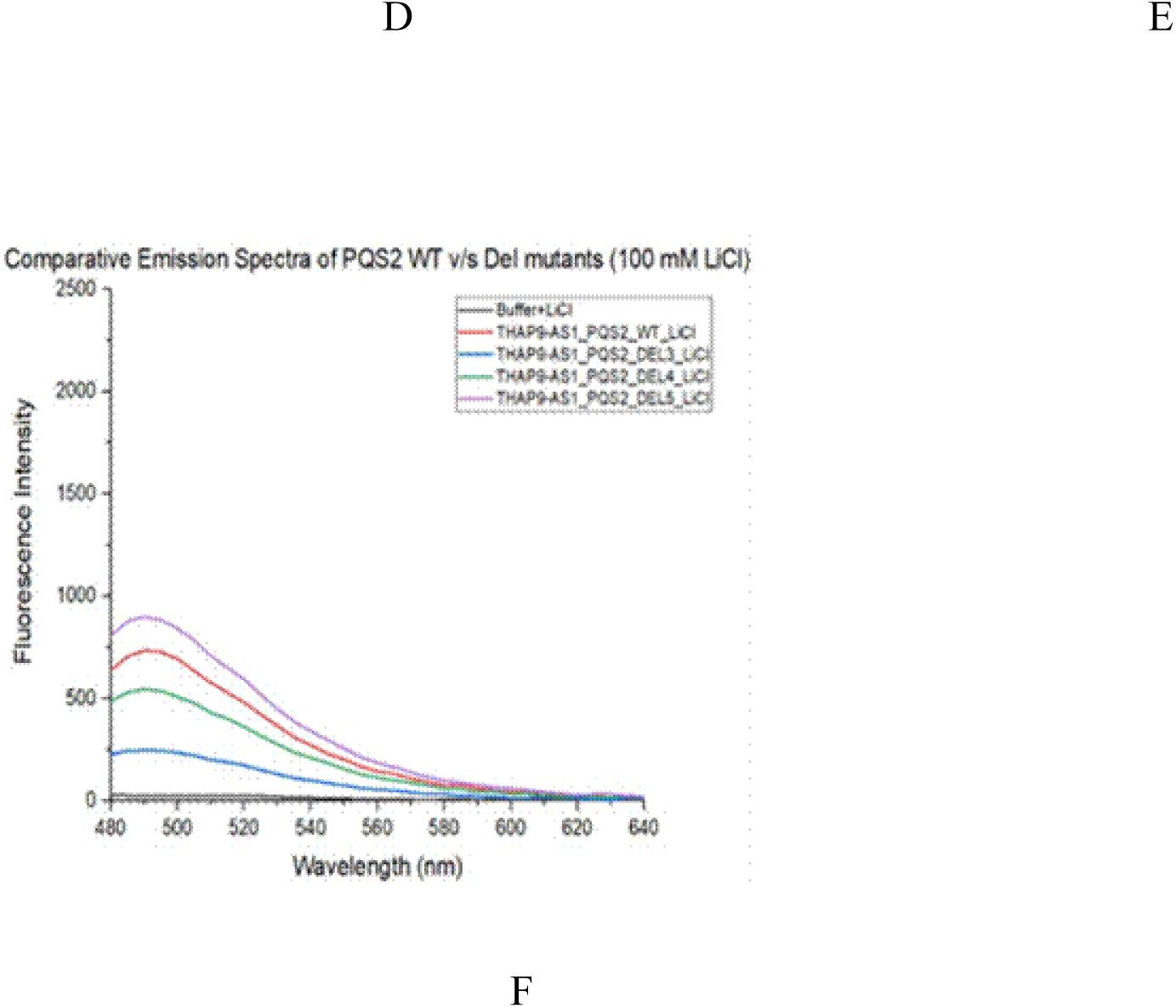
Emission spectra (when excited at 445 nm) of PQS1 and Deletion mutants (A) in the absence of K^+^ and Li^+^ (B) in the presence of K^+^, (C) in the presence of Li^+^. PQS2 and Deletion mutants (D) in the absence of K^+^ and Li^+^ (E) in the presence of K^+^, (F) in the presence of Li^+^ and ThT (2 µM)

**Figure 5.**
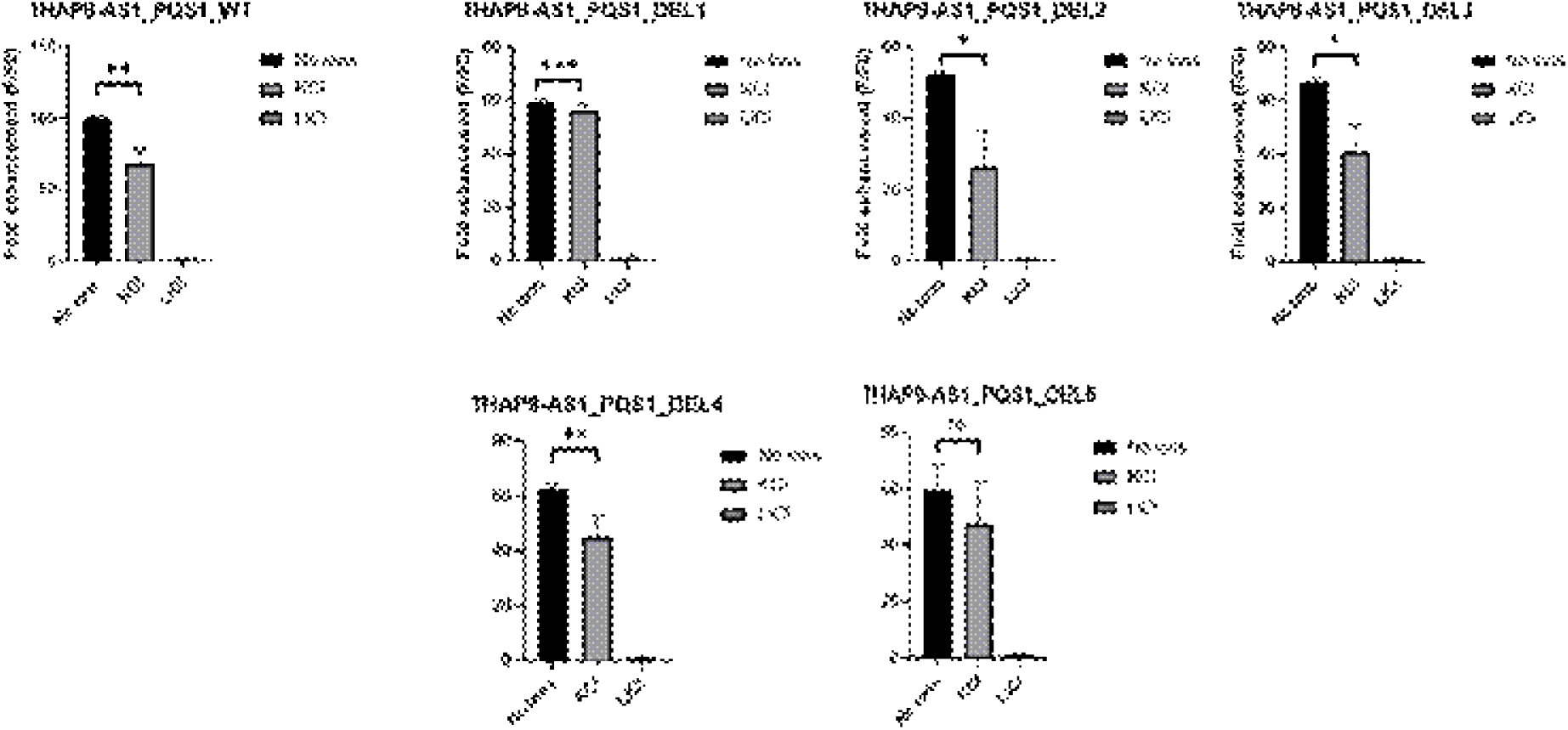
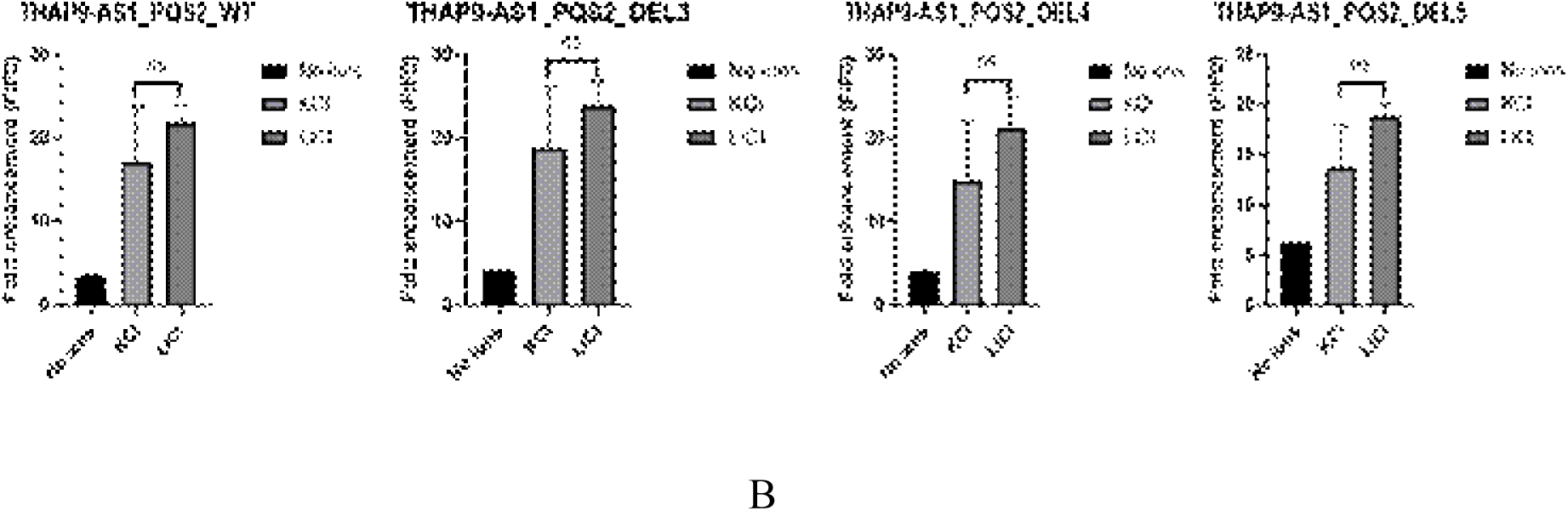
Fold enhancement of ThT fluorescence in the presence and absence of monovalent cations, with excitation and emission at 445 and 488, respectively, for (A) PQS1 and Deletion mutants (B) PQS2 and Deletion mutants. Data expressed as mean ± SEM. * p ≤ 0.05; ** p ≤ 0.001, *** p ≤ 0.0001.

PQS2-wild type shows higher fluorescence intensity than its deletion mutants in the absence of stabilizing monovalent ions (Fig. 4 (D)). On the other hand, for PQS2 deletion mutants, both K^+^ and Li^+^ ions have stabilizing effects on G4 formation (Fig. 5 (B)), as compared to absence of ions with Li^+^ ions having relatively more stabilizing effect (10-fold enhancement of ThT fluorescence with respect to presence of K^+^ ions). Moreover, PQS2 deletion mutant 5 (purple) is relatively more stable than even the wild type (red) in the presence of stabilizing monovalent ions like K^+^ (Fig. 4 (E)) and Li^+^ (Fig. 5 (F)).

Thus, it is observed that the enhancement of ThT fluorescence possibly induced by its binding to rG4 structure in the PQSs, is pronounced in the absence of monovalent ions but weakened in the presence of the monovalent ions (K^+^/Li^+^) in the case of PQS1. On the other hand, for PQS2, the effect of ions on the ThT fluorescence enhancement is reversed i.e. the effect is weakened in the absence but pronounced in the presence of the monovalent ions (K^+^/Li^+^). This phenomenon could be attributed to the stabilizing effect of certain monovalent cations on rG4 formation via 2G tracts in the case of PQS2. On the other hand, the presence of a 4G tract in PQS1 may allow it to form a stable rG4 structure even in absence of monovalent cations.

#### 2.2.5 Reverse Transcriptase stop assay

The ThT fluorescence assay is an indirect measure of the presence of rG4s as well as their stability since G4 formation is measured by the increase in the fluorescence intensity of the ThT molecule when it binds to the G4 structure. In the reverse transcriptase stop assay (RT stop assay), RNA is reverse transcribed by the reverse transcriptase enzyme until it is hindered by the presence of a G4 structure. Since, the assay is dependent on the physical presence of an rG4 structure and not just the downstream effects of its presence, it is a direct assay to determine the presence and stability of an rG4 structure in a sequence. Truncated complementary DNA products thus formed are visualized by running on a denaturing PAGE.

Thus, to confirm the presence of rG4s in PQS1 and PQS2 identified in THAP9-AS1, Reverse Stop assays were performed in the presence of increasing concentrations of monovalent cations, KCl and LiCl (highest concentration 150mM).

For both PQS1(Fig. 6 (A), (B)) and PQS2 (Figure 6 (C) and (D)), the denaturing gels illustrate that in the absence of monovalent cations, the band for the full-length product (Fig. 7 (A), (E)) had a much higher intensity than the stop product (Fig. 7 (B), (F)) which was not affected by increasing concentrations of K^+^. However, the presence of Li^+^ causes a marked decrease in the intensity of the full-length product band (Fig. 7 (C), (G)) and an increase in the stop product band intensity (Fig. 7 (D), (H)) with complete disruption of the full-length band at the highest concentration (150mM) of LiCl (Lanes 4 of Fig. 6 B and D). This suggests that for both PQS1 and PQS2, the stabilizing effect of Li^+^ is lower than that of K^+^.

**Figure 6.**
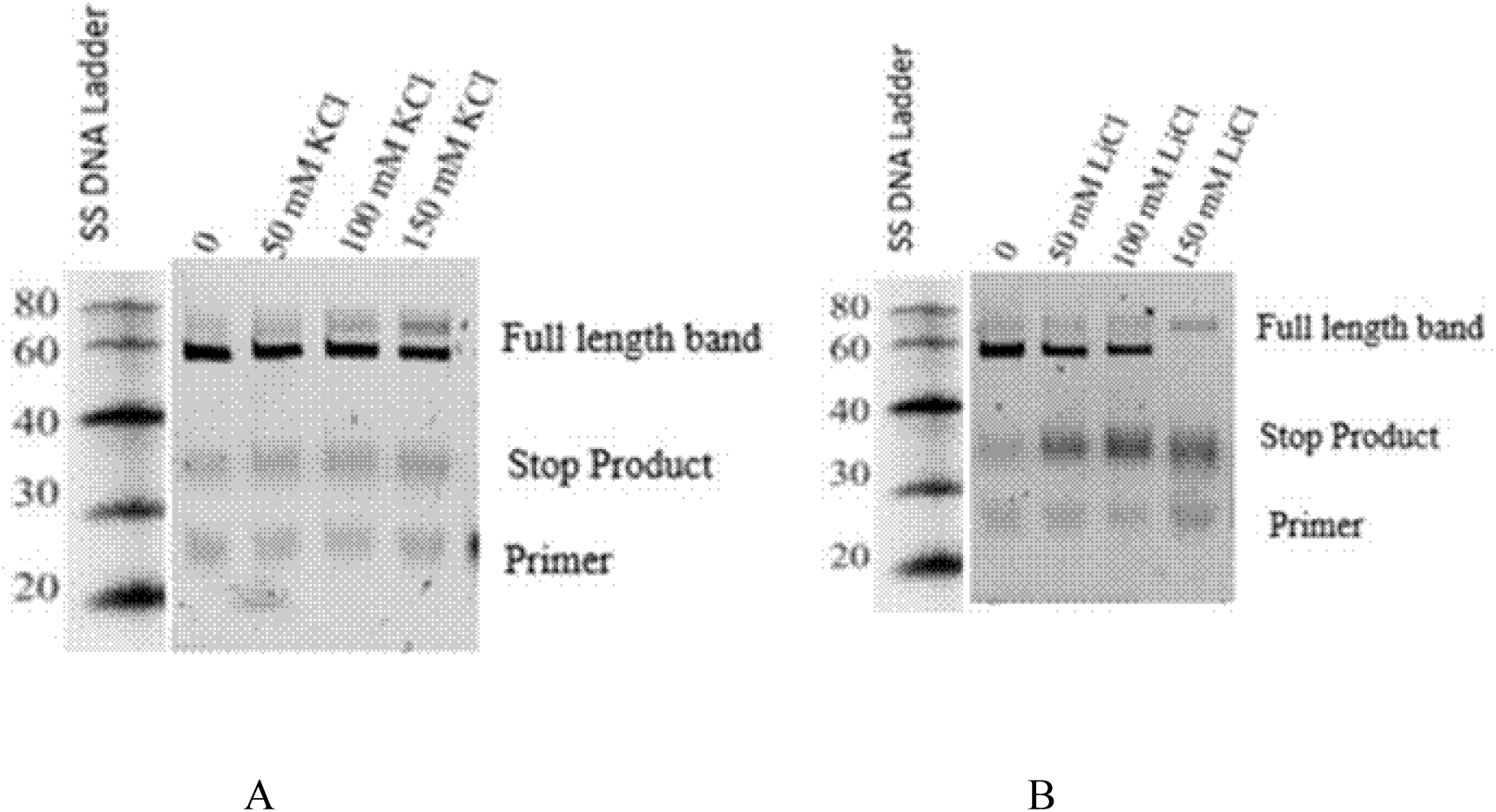
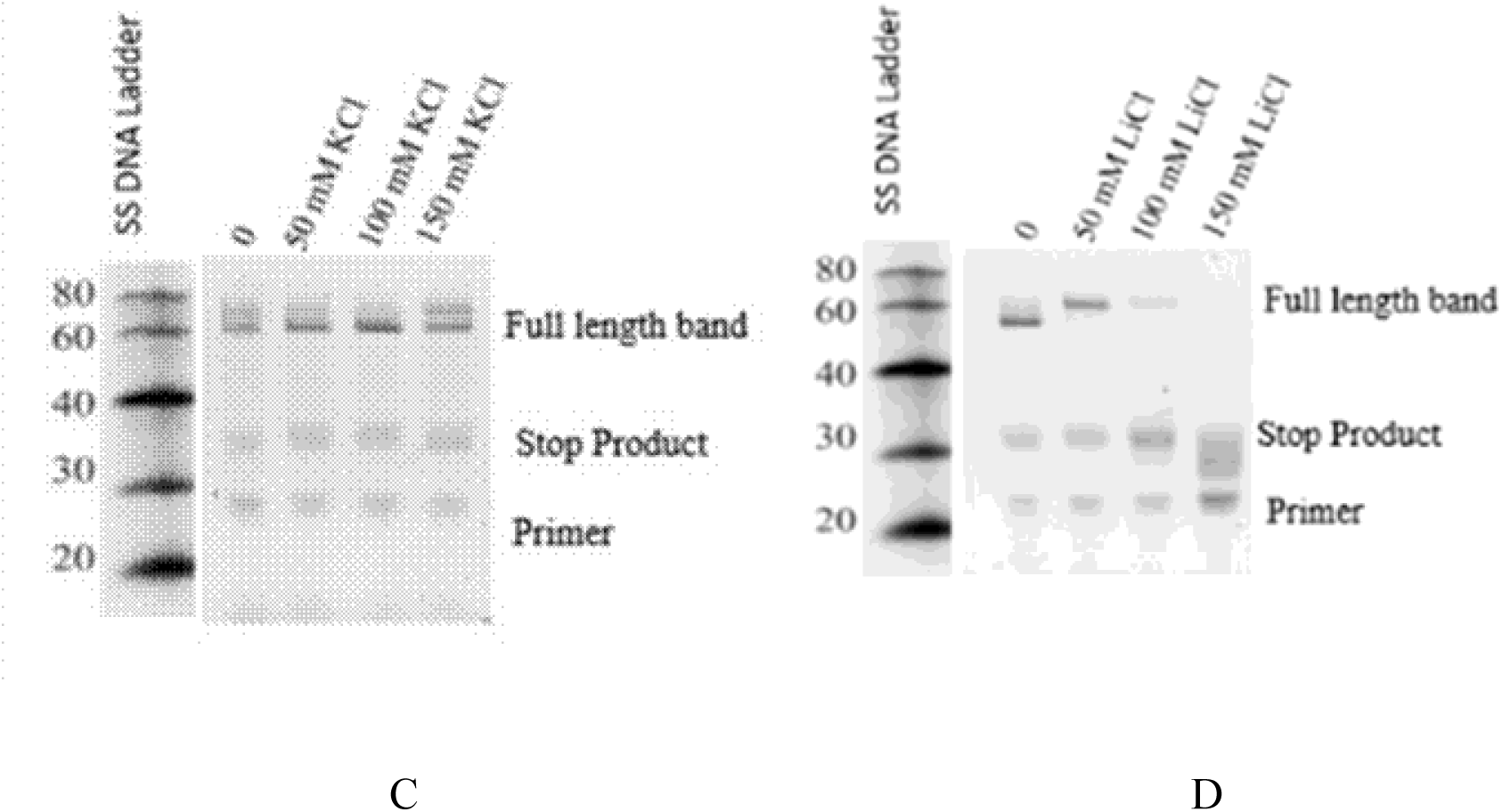
Denaturing Urea Gel of Reverse Transcriptase stop assay in the presence of increasing concentrations (0 -150 mM) of monovalent cations, for PQS1 (A) K^+^ (B) Li^+^ and PQS2 (C) K^+^ (D) Li^+^

**Figure 7.**
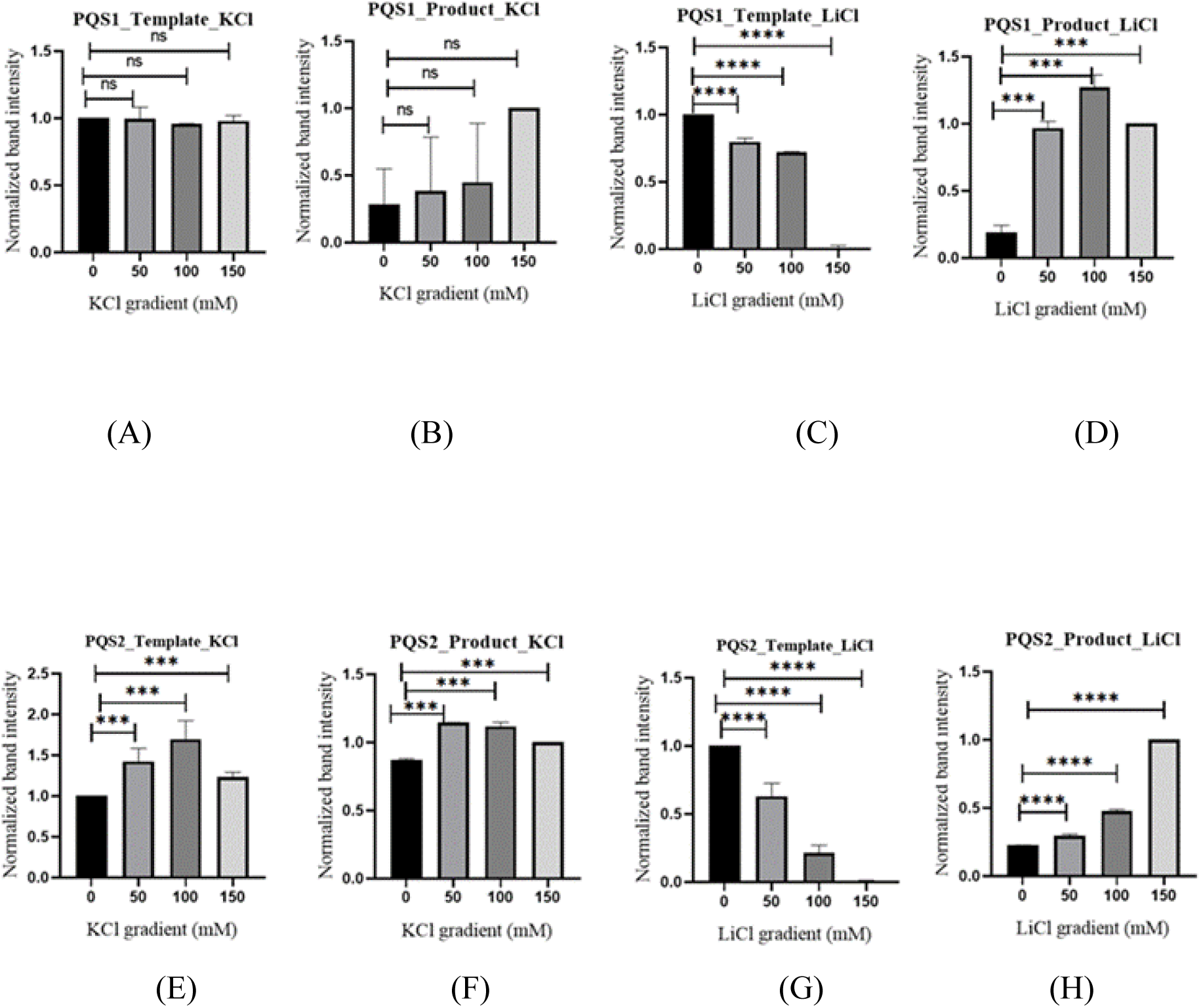
Quantification of full-length template bands and stop products produced in RT Stop assay (Figure 2.16). (A) PQS1 template in presence of KCl (0-150mM) (B) PQS1 product in presence of KCl (0-150mM) (C) PQS1 template in presence of LiCl (0-150mM) (D) PQS1 product in presence of LiCl (0-150mM) (E) PQS2 template in presence of KCl (0-150mM) (F) PQS2 product in presence of KCl (0-150mM) (G) PQS2 template in presence of LiCl (0-150mM) (H) PQS2 product in presence of LiCl (0-150mM) Ordinary one-way ANOVA was employed for the statistical analysis using GraphPad, and the resulting statistical significance is denoted with asterisks (*). Nonsignificant *P*-values are represented as ns.

KCl stabilizes the rG4 formed by the 2G tracts present in PQS2 and hence, the intensity of the template band increases at 50mM and 100 mM KCl but the rG4 is disrupted at higher KCl concentrations. LiCl disrupts the rG4 formed by PQS2 even at 50mM concentration, with an increase in stop product band intensity.

Thus, the Reverse transcriptase stop assay demonstrates that the stability of the rG4 is maintained in the presence of increasing concentrations of K^+^ ions while Li^+^ ions disrupt the rG4 structure.

Thus diverse experimental methods like Native Gel imaging, CD spectroscopy, ThT fluorescence enhancement and Reverse transcriptase stop assays demonstrate that the identified PQSs in THAP9-AS1 lncRNA exhibit the characteristic traits associated with G4 forming sequences. Native gel imaging illustrates G quadruplex formation via ThT staining. This is further validated by the presence of characteristic CD maxima at 265 nm and minima at 240 nm which is an indication of parallel G4 formation. The CD results are further substantiated with ThT fluorescence assay wherein the observed significant increase in fluorescence intensity is a result of ThT binding to the formed G4. The Reverse transcriptase stop assay which directly probes G4 formation demonstrates that both PQS1 and PQS2 form stable G4s with distinct stability response in presence of increasing concentrations (0 -150 mM) of monovalent cation K^+^ and Li^+^. These experiments successfully validate the G4 structure formation by the PQS identified via *in silico* methods.

The THAP9-AS1 lncRNA, which is implicated in various diseases (Su et al., 2022), is antisense to the THAP9 gene. The THAP9 gene codes for a homolog of the *Drosophila* P-element transposase. Interestingly, the THAP9-AS1 and THAP9 genes occur in a head-to-head fashion and share an overlapping promoter (Rashmi & Majumdar, 2022). Since sense and antisense genes often regulate each other (Villegas & Zaphiropoulos, 2015), it is interesting to hypothesise that THAP9 and THAP9AS1 may also regulate each other.

rG4s are often involved in interactions with proteins. This study was based on the hypothesis that THAP9 and THAP9AS1 may regulate each other. Towards this, we investigated if the rG4s in THAP9-AS1 lncRNA can interact with THAP9 since the latter contains a characteristic amino-terminal THAP domain that can bind nucleic acids like DNA through zinc-dependent mechanisms.

#### 2.2.6 RPISeq predicted binding potential of lncRNAs to THAP9 protein

The RPIseq bioinformatic prediction tool was used to investigate the interaction potential of THAP9-AS1 with THAP9 (Table 3). The interaction probabilities that are generated from RPISeq both for SVM and RF range between 0 to 1. The probabilities greater than 0.5 are considered to be positive i.e., the RNA is likely to interact with the protein. As can be observed from Table 3, both SVM and RF show values that vouch strongly in favour of the interaction of lncRNA THAP9-AS1 and full length THAP9 protein.

**Table 3.** RPISeq prediction of THAP9-AS1lncRNA interaction with THAP9 protein.

| **Sr. No.** | **LncRNA** | **Protein** | **Protein Length** | **SVM** | **RF** |
| --- | --- | --- | --- | --- | --- |
| 1. | THAP9-AS1 (Transcript variant 1) | THAP9 (Q9H5L6) | 903 a.a | 0.99 | 0.6 |
| 2. | THAP9-AS1 (Transcript variant 2) | THAP9 (Q9H5L6) | 903 a.a | 0.99 | 0.6 |
| 3. | THAP9-AS1 (Transcript variant 3) | THAP9 (Q9H5L6) | 903 a.a | 0.99 | 0.6 |

##### RPISeq Portal Data

Further, in Table 4, the potential interaction between the PQS1 and PQS2 (identified and validated in the previous section of the study) of THAP9-AS1 with the full length THAP9 is explored. The SVM and RF values suggest that the rG4s in THAP9-AS1, namely PQS1 and PQS2, also strongly bind the full length THAP9 protein.

**Table 4.** RPISeq prediction of PQS1 and PQS2 interaction with THAP9 protein (full length) and THAP domain.

| Sr. No. | LncRNA | Protein | Protein Length | SVM | RF |
| --- | --- | --- | --- | --- | --- |
| 1. | PQS1 | THAP9 (Q9H5L6) | 903 a.a | 0.91 | 0.55 |
| 2. | PQS2 | THAP9 (Q9H5L6) | 903 a.a | 0.78 | 0.7 |

#### 2.2.7 ITC

To investigate if the rG4s (PQS1 and PQS2) in THAP9-AS1 lncRNA can interact with THAP9, ITC was performed using each PQS (synthesised by *in vitro* transcription described before) and the recombinantly expressed THAP domain of THAP9. The full length THAP9 protein could not be included in the study since it did not express optimally.

The ITC results (Fig. 8) indicate a thermodynamically favourable interaction between the THAP domain and THAP9-AS1 lncRNA PQS1 and PQS2 (Table 5) with Δ*G* values of -80 Kcal/mol and -16.4 Kcal/mol and *K* _D_ values of 88.6 × 10^-9^ M and 199 × 10^-6^ M respectively. ΔG (Gibbs free energy change) represents the thermodynamic favourability and speed of binding; negative ΔG is indicative of a spontaneous binding interaction. The difference in the Δ*G* values and the *K*_D_ values between PQS1 and PQS2 hint at differences in the strength of their interaction with the THAP domain. This could be attributed to the presence of 4G tracts in PQS1 whereas PQS2 is made up of 2G tracts only.

**Figure 8.**
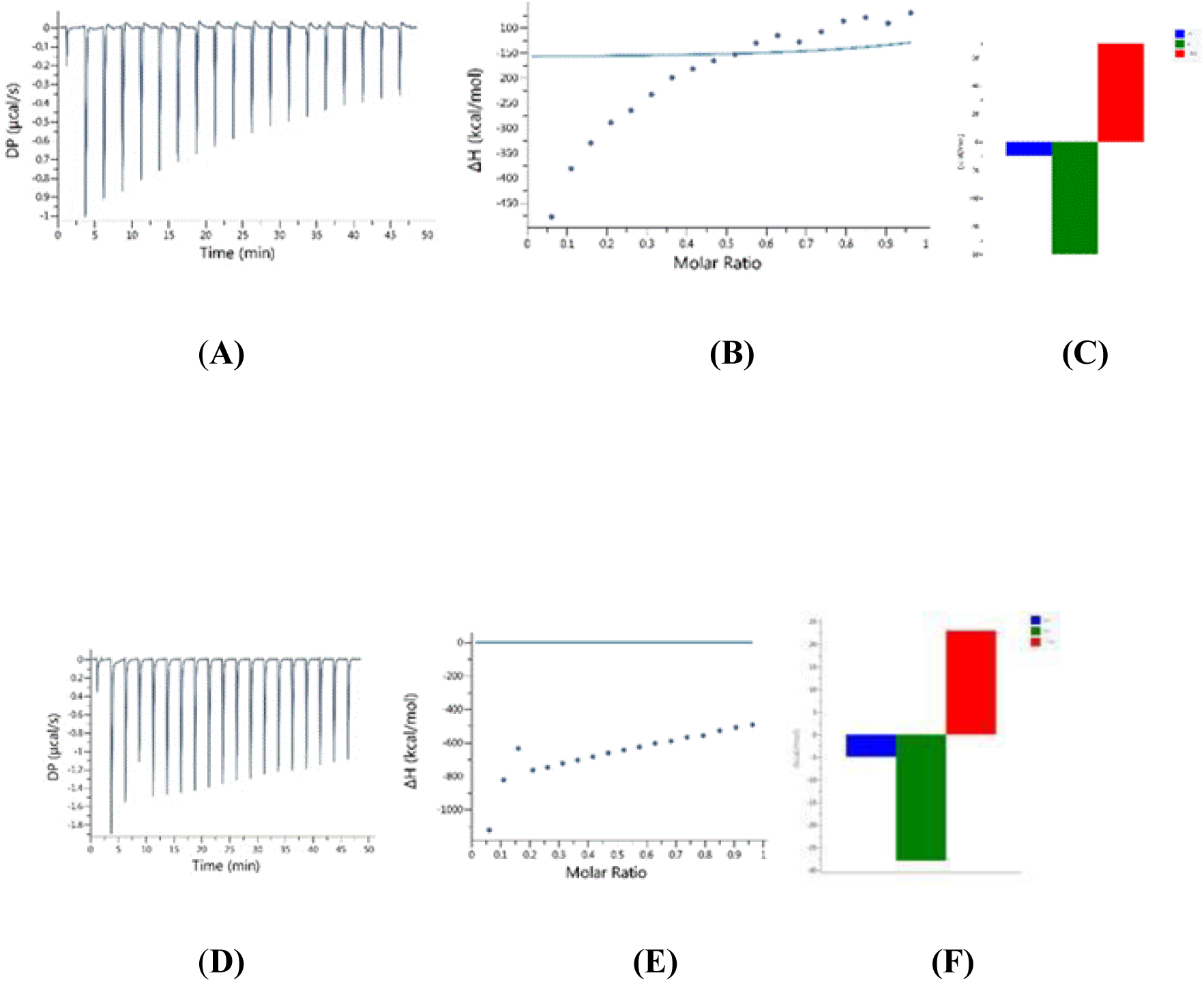
ITC of PQS1 and PQS2 with THAP Domain. A-C: PQS1, D-F: PQS2 (A) and (D) represent the raw ITC thermogram, wherein the X-axis is Time (min) showing injection of protein at fixed intervals of time leading to the spikes. The Y-axis represents DP (μcal/s). This represents the differential power applied by the instrument to keep the sample cell and reference cell at the same temperature. Each peak corresponds to heat released on binding. (B) and (E) represent the binding isotherm derived from the ITC thermogram. The X-axis represents molar ratio while Y-axis represents ΔH (Kcal/mol). The graph represents the reduction in free binding sites as the molar ratio of the titrant increases stating that the RNA is reaching saturation. In (C) and (F) the bar plot is a comparison of three thermodynamic properties. The blue bar represents ΔH, the green bar represents TΔS and the red bar represents ΔG. The binding enthalpy is weak however, the free energy change is significant and the negative entropy shows that the interaction is strongly entropy driven.

**Table 5.** Table representing the Δ*G* values and the *K* _D_ values obtained from ITC.

| Sequence name | $\Delta G$ (Kcal/mol) | $K_D$ (M) |
| --- | --- | --- |
| PQS1WT | -80 | $88.6 \times 10^{-9}$ |
| PQS2WT | -16.4 | $199 \times 10^{-6}$ |
| PQS2Δ3 | -13.6 | $664 \times 10^{-6}$ |
| PQS2Δ4 | -13.6 | $597 \times 10^{-6}$ |
| PQS2Δ5 | -13.6 | $84.2 \times 10^{-6}$ |

To investigate if the difference in binding affinities of PQS1 and PQS2 is indeed due to the presence of different G tracts, individual G tracts were systematically deleted. Subsequently, ITC-based binding studies (Fig. 8) were performed using deletion mutants of THAP9-AS1 PQS1 and PQS2. None of the deletion mutants of 4G tract-rich PQS1 showed binding (data not shown) with the THAP domain. The deletion mutants 1 and 2 of PQS2 are not used in this study as the deletion of first and second G-tract leads to complete loss of G4 formation as validated.

On the other hand, PQS2Δ3, PQS2Δ4, and PQS2Δ5 mutants showed significant binding with the THAP domain (Table 5, Fig. 9). It is observed that these mutants bind protein with varying strengths with Δ*G* values of -13.6 Kcal/mol and *K*_D_ values of 664 × 10^-6^ M, 597 × 10^-6^ M, and 84.2 × 10^-6^ M respectively (Table 5). While PQS2Δ3 and PQS2Δ4 may have weaker binding due to loss of G tracts, it is interesting to note that the binding by PQS2Δ5 may be slightly stronger than PQS2.

**Figure 9.**
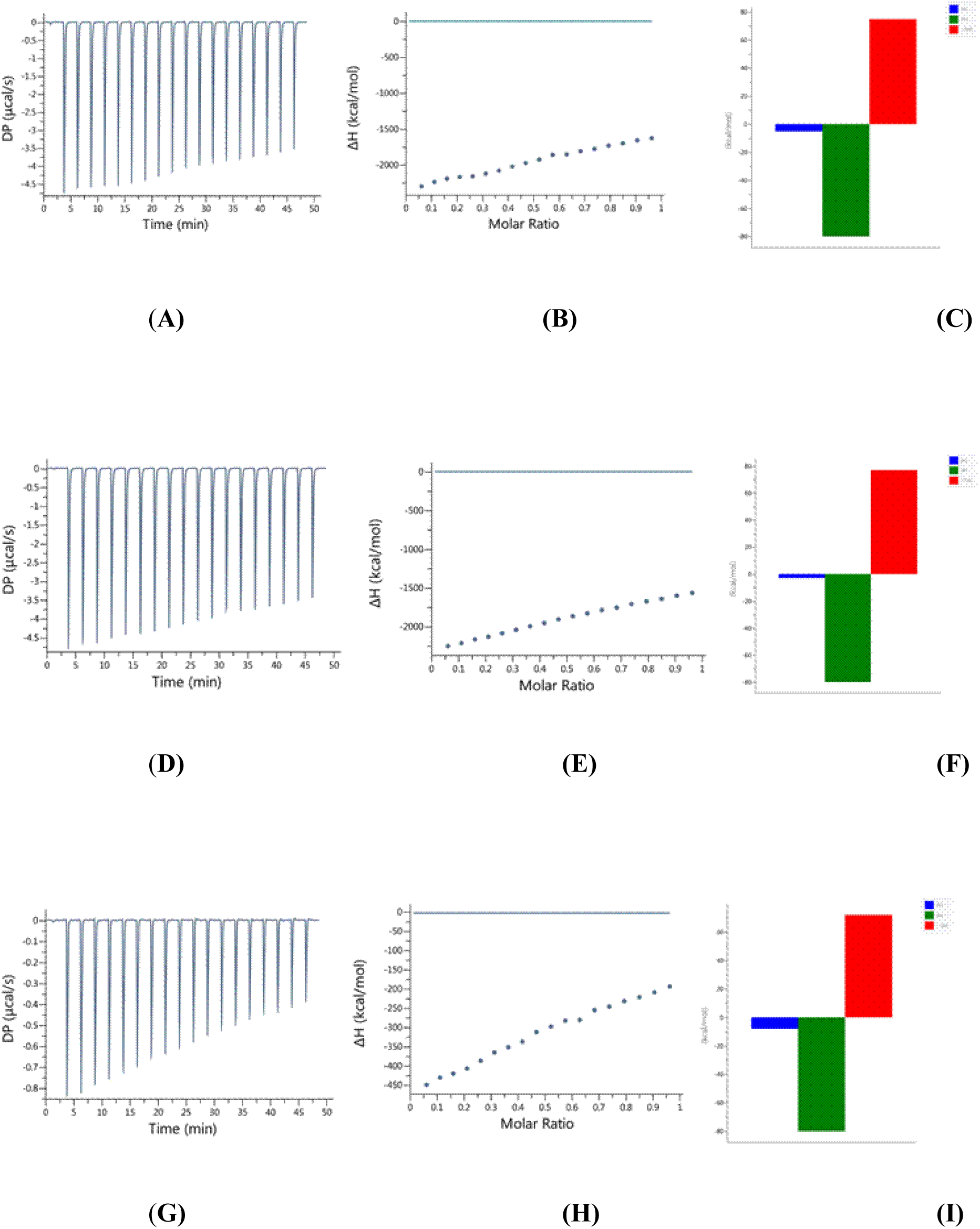
ITC of PQS2 deletion mutants with THAP Domain. A-C: PQS2 Deletion 3, D-F: PQS2 Deletion 4, G-I: PQS2 Deletion 5. (A), (D) and (G) Raw ITC thermogram, X-axis is Time (min), Y-axis represents DP (μcal/s). (B), (E) and (H) represent the binding isotherm derived from the ITC thermogram. The X-axis represents molar ratio while Y-axis represents ΔH (Kcal/mol). (C), (F) and (I) Bar plot comparing ΔH (blue bar), TΔS (green bar), ΔG (red bar).

Overall the ITC analysis demonstrates strong interaction between the THAP domain of THAP9 and both PQS1 and PQS2 of THAP9-AS1. However, these studies are unable to determine the exact bases of the RNA which bind the protein, whether this interaction is indeed rG4 mediated or whether the rG4 structure is unravelled or not during protein binding.

#### 2.2.8 Fluorescence spectroscopy

ThT fluorescence enhancement assay was performed to further investigate the interaction of the rG4s (in each of PQS1 and PQS2) with the THAP domain. The assay was performed with increasing concentration of THAP domain protein (5-30 μM). 40-fold decrease and 30-fold decrease in the ThT fluorescence was observed for PQS1 and PQS2 respectively, with increasing concentrations of THAP domain. The inversely proportional relation i.e., decrease in fluorescence with increasing protein concentration hints at two possible scenarios that could occur when PQS1 and PQS2 interact with the THAP domain. The THAP domain may competitively bind to the rG4 and inhibit the binding of ThT molecule to the rG4 structure. Alternatively, the binding of the THAP domain to the rG4 can alter the conformation of the rG4 such that ThT binding can no longer be facilitated. Either of the two scenarios could lead to reduction in the fluorescence in the ThT fluorescence enhancement assay. Both the PQS1 and PQS2 show similar change in pattern with respect to ThT fluorescence.

The ThT fluorescence assay shows a drastic attenuation in the fluorescence even at the lowest concentration of the protein (5 μM) (Fig. 10). This drastic reduction in ThT fluorescence hints at strong interaction between the THAP domain protein and rG4. The THAP domain could be hindering the binding of ThT with the rG4 structure when complexed with the RNA or modulating the rG4 structure when it binds to the RNA.

**Figure 10.**
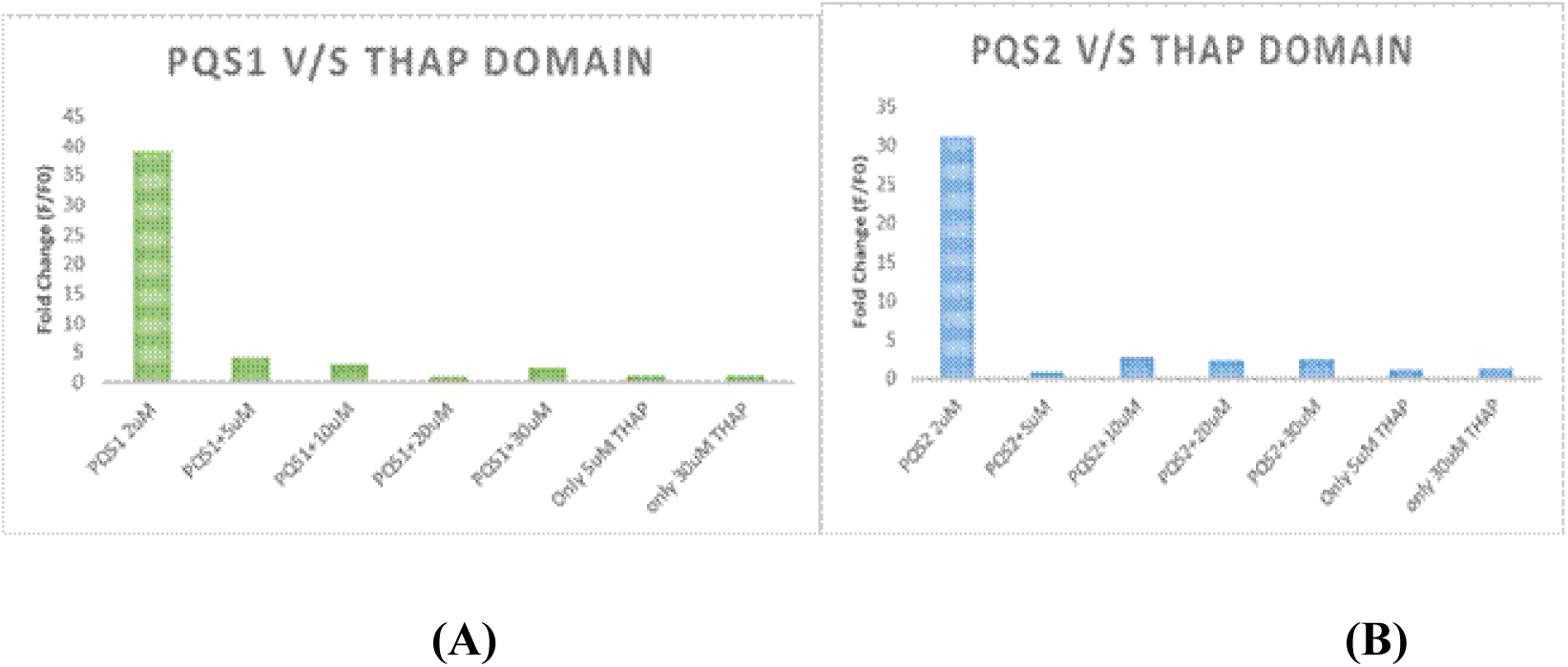
ThT fluorescence enhancement assay of lncRNA THAP9-AS1 PQS1 and PQS2. Fold enhancement of ThT fluorescence for G4 lncRNA THAP9-AS1 (A) PQS1 (2 μM) (B) PQS2 (2 μM) in the presence of THAP domain (5 μM to 30 μM). Excitation and emission were at 445 and 488 nm, respectively. An unpaired *t*-test was used for statistical analysis, and all the resulting *P*-values were significant (p ≤ 0.05)

#### 2.2.9 Electrophoretic Mobility Shift Assay

To further investigate the interaction of the rG4s formed by the PQS1 and PQS2 with the THAP domain protein, EMSA was performed. Although the ThT fluorescence assay sheds some light on the dynamics of the rG4 structure (protein binding could either unravel the rG4 or prevent ThT binding to the rG4) in both the PQSs when interacting with the THAP domain, it does not reveal much about the change in molecularity (i.e. number of interacting molecules of RNA and protein) of the interacting partners. It is possible that PQS1 and PQS2 could interact in different stoichiometries with the THAP domain. To ascertain the changes in molecularity, EMSA was performed.

2 μM of RNA was incubated with increasing concentrations of THAP domain protein (2 μM to 12 μM). The binding of the RNA to the protein should lead to a change in its electrophoretic mobility. The gels were imaged using a Rhodamine filter to detect the presence of primer bound RNA and then counterstained with ThT and visualized at 488 nm to detect rG4 formation. As controls, increasing concentrations of only wildtype PQS1 and PQS2 folded RNA were run on a gel and stained with ThT dye (Fig 11 (E)) to determine rG4 presence and Diamond Nucleic acid (Fig 11 (F)) to determine RNA abundance.

**Figure 11.**
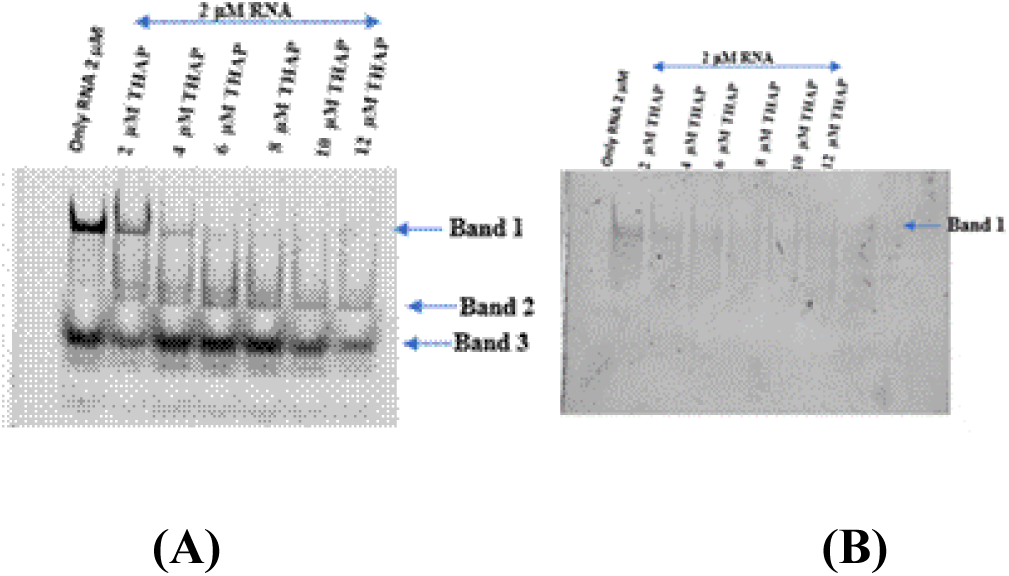
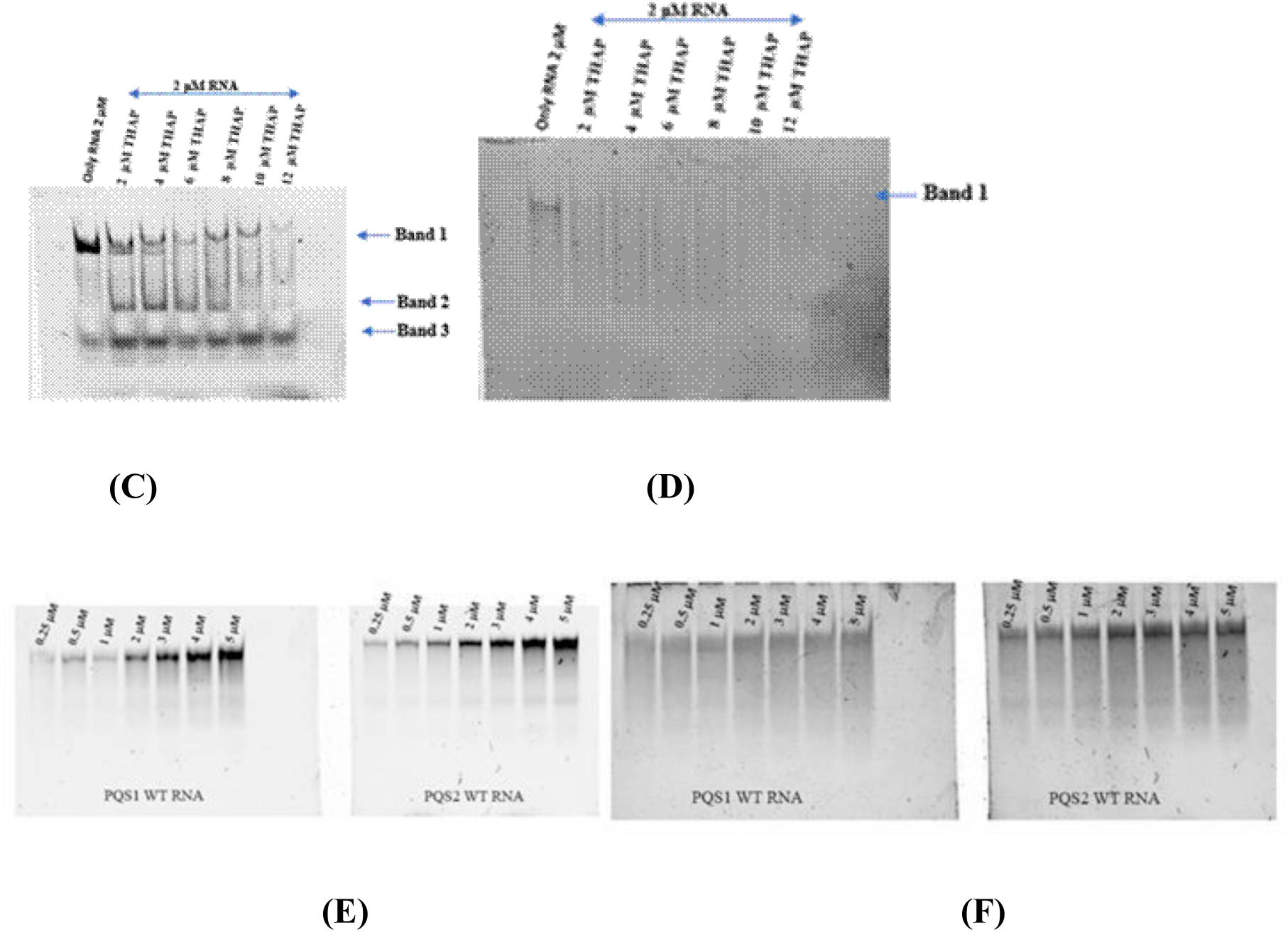
EMSA of lncRNA THAP9-AS1 PQS1 and PQS2 with THAP domain. Annealed and folded THAP9-AS1 PQS1 (2 μM) with increasing concentration of THAP domain (2 μM to 12 μM) (A) visualized in rhodamine filter. (B) with ThT. Annealed and folded THAP9-AS1 PQS2 (2 μM) with increasing concentration of THAP domain (2 μM to 12 μM) (C) visualized in rhodamine filter. (D) with ThT Only PQS1 and PQS2 WT folded RNA in increasing concentration stained with (E) ThT dye (F) Diamond Nucleic acid

Fig. 11 (A,C) illustrates the presence of multiple bands (Bands 1, 2, 3) when the gel is visualized using Rhodamine filter (detect primer bound RNA). Band 3 present across all lanes is the dye front. Band 1 corresponds to unbound lncRNA and is present in all lanes including the first lane that contains lncRNA alone, devoid of protein. In case of PQS1 (Fig. 11 (A)), the intensity of the rG4 band 1 decreases with increase in protein concentration, suggesting depletion due to binding with the THAP domain protein as suggested by the ITC results (Fig. 8). In case of PQS2 (Fig. 11 (C)), the RNA may be forming higher order intermolecular multimeric species as seen from the shift in band 1. Band 2 corresponds to the lncRNA + THAP domain complex.

Fig. 11 (A) and (C) illustrate that with increase in protein concentration, the RNA binds strongly to the protein reducing the hydrodynamic radius of the complex, thereby allowing it to migrate faster through the gel. This could be attributed to protein induced changes in the RNA (Ryder et al., 2008). Traditionally, protein-nucleic acid complexes have slower electrophoretic mobility in EMSA; however, a change in the hydrodynamic radius and shape of the complex can sometimes lead to counterintuitive mobility shifts.

In case of both PQS1 and PQS2, counter-staining the gels with ThT gels shows no evidence of G4 structure retention after protein binding (Fig. 11 B, D). This suggests that the THAP domain protein causes disruption of the G4 structure or displacement of ThT from the G4 binding site. It can be ascertained that the observed variation in molecularity (Fig. 11 A-D) is a function of the protein binding to the RNA because the control gels (Fig. 11 E, F) illustrate that at higher concentrations of RNA (alone in absence of protein) there is no change in its molecularity.

In summary, we conclude that the rG4 of THAP9-AS1 lncRNA interacts strongly with the THAP domain of human THAP9 protein. This is corroborated by ITC analysis (higher *K*_D_ value, Table 5), EMSA analysis (Band 2 corresponding to RNA-protein complex forms at lower protein concentrations) as well as significant reduction in ThT fluorescence even at lowest concentration of protein (Fig. 10) which suggests a loss or occlusion of the rG4 structure due to protein binding. The lack of THAP domain binding to any of the deletion mutants of PQS1 hints at the importance of all the G-tracts in facilitating the lncRNA protein interaction.

The binding of PQS2 to the THAP domain is relatively weaker than PQS1, as can be inferred from the corresponding *K*_D_ value (Table 5). However, the significant reduction in ThT fluorescence at lowest protein concentration (Fig. 10), as well as the formation of higher molecular weight complexes in Band 2 at higher protein concentrations suggests that the THAP domain does bind to PQS2 (Fig. 11 C). In case of PQS2 too, there is loss or occlusion of rG4 structure due to the protein interaction. However, unlike PQS1, in the case of PQS2 the deletion mutants 3, 4, and 5 are all able to interact with the THAP domain (Fig. 9). This suggests that even in the absence of certain G tracts, the other G tracts compensate and form the G4 structure which is sufficient to facilitate the interaction.

Taken together, these results illustrate that the rG4 structures in the PQS1 and PQS2 of THAP9-AS1 lncRNA are key elements in recognition of protein binding partners.

The interaction of THAP9-AS1 lncRNA PQS1 and PQS2 with the THAP domain of the THAP9 protein is worth further study so as to identify its potential role in the context of the cell.

## 3. Discussion

This study is based on the quest to identify non canonical G4 forming tendencies present in THAP9-AS1 lncRNA which would help the RNA to interact with its binding partners. *In silico* tools were used to identify G quadruplex forming sequences (PQS) that are present in all three known transcript variants of THAP9-AS1. The study experimentally validates the identified PQSs using a combination of techniques like CD spectroscopy, ThT fluorescence assay, reverse transcriptase stop assay and Competition Assay (shown in supplementary). It is to be noted that the study was conducted with *in vitro* transcribed (IVT) RNA which is made of both full length transcripts as well as truncated products.

Overall, we observe that PQS1 and PQS2 both adopt stable parallel rG4 structures both in the absence and the presence of certain monovalent cations (K^+^). Li^+^ ions disrupt the rG4 structure. Moreover, PQS1 is significantly more stable than PQS2.

Overall, we observe that PQS1 and PQS2 both adopt stable parallel rG4 structures both in the absence and the presence of certain monovalent cations (K^+^). Li^+^ ions disrupt the rG4 structure. Moreover, PQS1 is significantly more stable than PQS2.

The *in vitro* identification of stable G4 structures in THAP9-AS1 lncRNA suggests that it may play a role in helping the lncRNA sequester miRNA and interact with proteins and DNA.

rG4s are structural motifs present on the lncRNA that play significant roles in facilitating interactions with biomolecules in the cell. No systematic study on the rG4 forming potential of THAP9-AS1 has been conducted before and hence no studies exist on the importance of rG4 motif formation.

In this study, a blend of methods like bioinformatics, fluorescence spectroscopy and isothermal titrimetric calorimetry have been used to explore the binding potential of rG4 motifs present in PQS1 and PQS2 of THAP9-AS1 lncRNA to the THAP domain of THAP9 protein (product of its sense gene pair). The results suggest a strong binding potential of THAP9-AS1 lncRNA with the THAP9 protein.

It is to be noted that the assays have been conducted with *in vitro* transcribed RNA, which is typically heterogeneous and can contain some shorter products along with desired full length product. Future studies can be conducted with IVT synthesized HPLC (High Performance Liquid Chromatography) purified RNA which would only comprise the desired full length strand. This would help shed better light on the interaction between THAP domain of THAP9 protein and PQSs identified in lncRNA THAP9-AS1.

lncRNAs are known to have multivalent interactions with RBPs so as to form paraspeckles (Yamazaki et al., 2018) as well as act as nucleation centres for sequestering ribonucleoprotein complexes (Lee et al., 2016). The EMSA assay shows that RNA, in absence of protein, even at very high concentrations, does not show a change in molecularity. However, protein binding leads to the formation of multiple RNA bands at various positions in the gel, hinting at change in molecularity of the RNA due to binding of the THAP domain at multiple positions. This interaction of THAP9 protein with the rG4 is further validated via ThT assay where the quenching of ThT is observed when the RNA is incubated with even the lowest concentration of protein (5 μM).

It is to be noted that the experiments performed in this study to make the above inferences, are all indirect methods as they do not allow direct visualisation of the rG4 of PQS1 and PQS2 or their interaction with the THAP domain of THAP9 protein. In the future, there is a need to shed further light on the structural dynamism at play in the presence or absence of protein, using methods like NMR, X-Ray diffraction crystallography.

## 4. Methods

### 4.1 Bioinformatic prediction of PQS in THAP9-AS1

The quadruplex forming ability of THAP9-AS1 was predicted and PQSs were identified using various G4 finder tools like imGQfinder, G4 hunter, and QGRS mapper (accessed on May 18, 2023).

### 4.2 Oligonucleotides

DNA Oligonucleotides were designed for *in vitro* transcription using T7 RNA polymerase and ordered from Sigma. The high yield T7 RNA promoter sequence (5’ TAATACGACTCACTATAGCGAAA 3’) was ordered separately from Sigma (Table S3) and annealed to the antisense DNA sequence as per the protocol provided by Sigma-Aldrich, St. Louis, MO, USA (Rychlik et al., 1990).

The annealing mixture is heated at 95^◦^C for 5 min and cooled to room temperature. The concentration and purity of the annealed DNA was quantified using NanoDrop™ 2000 spectrophotometer (Thermo Fisher Scientific).

*In vitro* transcription was performed using HiScribe™ T7 High Yield RNA Synthesis Kit (New England Biolabs, Catalogue no. E2040S), following the manufacturer’s protocol (Fig. 10). Dithiothreitol (reducing agent) is added to the reaction mixture to stabilize the enzymes. The template for the IVT reaction (T7 promoter sequence annealed to the DNA oligonucleotide) was digested using DNase I, RNase-free (Thermo Fisher Scientific, Catalogue No. EN0521) at the end of the reaction. RNAs were cleaned and eluted using Monarch® RNA Cleanup Kit (New England Biolabs, Catalogue No. T2050L), following the manufacturer’s protocol. The purity and concentration of the RNA was measured using a NanoDrop™ 2000 spectrophotometer. The purified RNA was run on 15% native PAGE at 120V for 60 min. The gels were further stained with ThT to visualize rG4 structure formed, and stained with Diamond nucleic acid to check for RNA abundance. The stained gels were visualized using the ChemiDocTM MP Imaging system (BioRad) at 488 nm filter and EtBr (Ethidium Bromide) filter for ThT and Diamond nucleic acid stains respectively.The purified RNA was stored at −80^◦^C.

### 4.3 CD Spectroscopy

Circular Dichroism (CD) spectroscopy was performed to evaluate the conformation of G-quadruplexes in RNA. CD spectra were recorded on a JASCO J-815 spectrophotometer at 16^◦^C in the wavelength range of 220–320 nm, using a response time of 1 s, a step size of 1 nm and a 1 nm bandwidth. The scanning speed of the instrument was set at 100 nm/min, with an average of three scans. A 1 mm path length quartz cuvette was used in all experiments. Samples containing 5 µM RNA were folded in a buffer containing 10 mM Tris-Cl (pH 7.5), 0.1 mM EDTA (pH 8.0) by incubating at 95°C for 5 min and cooled to room temperature before CD analysis.

### 4.4 Fluorescence spectroscopy

The Fluorescence enhancement assay was performed using the RNA G4 binding dye, Thioflavin T (ThT, Sigma-Aldrich, USA, Catalogue no. 2390-54-7) in a 384-well black fluorescence microplate using a Fluorescence Microplate Reader (Synergy H1M, M/s Biotek Instruments Inc. USA). ThT is a benzothiazole dye which when dissolved in a buffer consisting of 10mM Tris pH 7.5 and 100uM EDTA exhibits negligible fluorescence emission at 488 nm. However, ThT undergoes multiple fold increase in fluorescence when it binds to a G4 structure and its rotational freedom is restricted. The planar aromatic ring of ThT is stacked on the exposed surface of the G quartet. The π-surface of the G quartet facilitates π–π interactions with the ThT molecule. This interaction immobilizes the C-C bond causing strong fluorescence enhancement (De La Faverie et al., 2014; Mohanty et al., 2013).

The RNA sample (2 µM) was folded in a buffer consisting of 10 mM Tris-HCl (pH 7.5) and 0.1 mM EDTA (pH 8.0) supplemented with monovalent cations (K+/Li+ 100mM). The mixture was then heated at 95°C for five minutes and cooled to room temperature gradually. ThT (2 µM) was added to the folded RNA and excitation spectra were recorded with emission at 488 nm. An emission spectra was also obtained after excitation at 445 nm. Single point fluorescence intensities were also captured for ThT at the mentioned wavelengths. All spectra were collected at 37°C.

### 4.5 Reverse Transcriptase stop assay stop assay

Texas red labelled primers (length=20 nucleotides, 5’ TGTGTTCTTGTTCTCTGTCG 3’ which binds to Random primer binding site at 3’ end of PQS RNA; Fig 9]) were purchased from Sigma Aldrich, USA in lyophilized form and nuclease-free water was used to prepare 100 µM solutions. Each reverse transcriptase stop assay was performed in a 10 µL reaction mixture (Table 2.2) consisting of 1X Annealing buffer (pH 7.5) 2 µM RNA, 100 nM Texas Red labelled primers, 2 mM NTPs and monovalent cations (150mM KCl/LiCl). The RNA is annealed to the labelled primers by heating to 95°C for five minutes and then cooled to room temperature gradually. Reverse transcriptase enzyme (M-MLV Reverse Transcriptase RNase H minus, Promega, catalogue no., M5301) is then added to the reaction and incubated for 1 h at 37 °C. At the end of the incubation period, the reaction was terminated by addition of a buffer containing 95% Formamide, 0.05% Bromophenol Blue, 20 mM EDTA (pH 8), and 0.05% Xylene cyanol. The reaction products are then resolved using 15% denaturing (Urea) PAGE. The gel is visualized on a ChemiDocTM MP Imaging system (BioRad) using the Rhodamine filter and counter stained with DiamondTM Nucleic acid dye (Promega, Catalogue no. H118A) to observe template bands.

The identified PQS. PQS1 and PQS2 are further subjected to bioinformatic analysis to predict their binding potential to THAP9 protein (product of the sense gene pair THAP9).

### 4.6 Bioinformatic approach to investigate binding potential of THAP9-AS1 to THAP9

RPISeq is an online RNA-Protein Interaction prediction tool which predicts if an RNA and protein molecule will interact based on the sequence of the RNA and protein. The RNA and protein sequences are taken as input in the online tool which then gives interaction probabilities values generated via SVM (Support Vector Machine) and RF (Random Forest) classifiers (Muppirala et al., 2011).

### 4.7 THAP domain of human THAP9

The cDNA of hTHAP9 N-terminal THAP domain (1-89 amino acid residues) with a C-terminal 6X histidine tag has previously been cloned into the pRSETa vector downstream of the Lac promoter. The vector was transformed into BL21DE3 cells for heterologous protein expression. The transformed cells were grown in Luria Bertani broth (Catalogue No. G1245 Himedia^TM^) overnight at 37°C as primary culture. This primary culture was further used as inoculum for secondary culture grown at 37°C. The secondary culture was induced with 1mM IPTG (TCI^TM^ CAS No. 367-93-1) once the OD reached 0.5. Post induction, the cells were grown at 37°C for 12 hours. Post 12 hours the cells were pelleted down at 6000 rpm for 15 minutes at 4°C. To purify the protein, the cells were resuspended in lysis buffer (20mM HEPES-KOH, pH-7.4, 500mM NaCl, 5mM MgCl2, 20mM Imidazole, glycerol 10% (v/v) 1mM PMSF), and subjected to lysis via sonication (ON- 15secs, OFF- 45secs, Amplitude- 50% using a VibraCell^TM^ Sonicator). The cell lysate was then pelleted via centrifugation at 40,000 rpm for 45 minutes at 4°C, using ultracentrifuge (Beckman Coulter^TM^). The supernatant was then incubated with 1mL Nickel-Sepharose High performance beads as described by manufacturer (GE Healthcare) for 3 hours at 4°C, followed by elution using imidazole-containing buffer. The THAP domain protein eluted at 200-300 mM imidazole concentration. The eluted protein was visualized via SDS-PAGE and was further concentrated in 3Kda centrifugal filters (Merck) as per the instruction manual. The concentrated proteins were then further subjected to size exclusion chromatography in AKTA-FPLC with Superdex 75 10/300GL column so as to get the desired pure 10kDa protein. The eluted proteins were then subjected to SDS-PAGE so as to confirm their purity and their purity was further validated using theoretical molar extinction coefficient on Nanodrop (Thermoscientific^TM^). The protein was stored at -80°C in 10% (v/v) glycerol.The protein was later subjected to buffer exchange into 1X Annealing buffer (10 mM Tris-Cl pH 7.5 and 0.1 mM EDTA pH 8.0) for the binding studies with RNA.

### 4.8 Fluorescence spectroscopy

Fluorescence enhancement assays were performed using Thioflavin T (ThT) (Catalog no. T3516, Sigma-Aldrich, USA) in a 384-well black fluorescence microplate. RNA samples (2 μM) were folded in a folding buffer containing 10 mM Tris-Cl pH 7.5 and 0.1 mM EDTA pH 8.0 by incubating at 95 °C for 5 min, followed by gradually cooling to room temperature. ThT (2 μM) was added to the folded RNA, and the emission spectra were obtained after excitation at 445 nm. The excitation spectra were obtained with emission capture at 488 nm. Single-point fluorescence intensities were also obtained for ThT at the mentioned wavelengths. The fluorescence of samples was measured at 25 °C using a Cytation 5 Cell Imaging Multimode Reader (Agilent Technologies, USA). The mean fluorescence intensities (in arbitrary units) were extracted from the emission spectra and were plotted against the wavelength after applying the Savitzky–Golay smoothing method with a 20-point window using Origin (Pro) software. For the ThT Fluorescence enhancement assays with Human THAP domain protein, the RNA was first folded as described and then incubated with the indicated concentrations of THAP domain for 2 h at 4 °C. Fluorescence measurements were performed as described above. An unpaired *t*-test was conducted for statistical analysis.

### 4.9 Isothermal Titrimetric Calorimetry (ITC)

ITC measures the enthalpy change (ΔH), which is the measure of heat absorbed or released on formation of a complex. It gives an idea regarding strength and nature of intermolecular interactions. Gibbs free energy (ΔG) represents the thermodynamic favourability and the speed of the binding. The more negative the ΔG the more spontaneous the binding reaction. TΔS refers to the change in entropy. It is calculated by using the formula TΔS = ΔH – ΔG. TΔS is a reflection of the changes in conformation and degrees of freedom associated with the binding.

ITC measurements were conducted using an ITC200 microcalorimeter (Microcal-200, Malvern Panalytical) at 25 °C. All lncRNA samples were folded in the folding buffer (10 mM Tris-Cl, pH 7.5, and 0.1 mM EDTA, pH 8.0) prior to titrations. The revolving syringe (750 rpm) was loaded with 10 μM Human THAP domain protein dissolved in the same buffer, and a 2μM lncRNA solution was injected inside the cell. Twenty injections of 40 μL of Human THAP domain protein were made into RNA, with a 150 s gap between each injection. To accommodate diffusion of the syringe into the cell during equilibration, the first injection was 0.4 μL, and data fitting did not include this injection. The isotherms were analyzed by using a single-site binding model in the Microcal ITC software to determine the relevant thermodynamic parameters.

### 4.10 Electrophoretic mobility shift assay (EMSA)

The primer binding site (described in section 2.5 and shown in figure 9) was used to label the synthesized lncRNAs with Texas-red-labeled primer for visualizing electrophoretic mobility shift assays. 2 μM RNA was first folded and annealed with the labeled primer by incubating at 95 °C for 5 min, followed by gradually cooling to room temperature. The primer-annealed RNA was then incubated with different concentrations of THAP domain protein for 2 h at 4 °C in 10 μL of total reaction volume. THAP domain protein solutions were prepared in a folding buffer (10 mM Tris-Cl, pH 7.5, and 0.1 mM EDTA, pH 8.0) and 5% glycerol. Binding reactions were loaded onto 15% polyacrylamide gels in 1× Tris-borate EDTA. The gels were imaged using a ChemiDoc MP Imaging system using the Rhodamine filter to detect the presence of primer bound RNA and then counterstained with 0.5 mM ThT to to detect the presence of G4 forming species. Band intensities were quantified using ImageJ software. Ordinary one-way ANOVA was employed for statistical analysis.

## 5. Conclusion

In this study, we have used *in silico* techniques to identify G4 forming sequences in THAP9-AS1 lncRNA. We selected PQS1 and PQS2, which are present in all 3 transcript variants of THAP9-AS1 and have experimentally characterized their G4 formation propensity by employing different approaches. The study demonstrates the presence of non-canonical G4 forming sequences in the lncRNA THAP9-AS1 thus validating the in silico predictions.

The study further reinforces the functional significance of rG4 present in PQS1 and PQS2 thereby facilitating molecular recognition and protein interaction.

This is the first reported biochemical investigation of the structural properties and G-quadruplex forming potential of a newly annotated lncRNA THAP9-AS1 that is implicated in diseases like cancer. There are no previously reported studies that show a lncRNA binding to the product of its sense strand gene pair. Thus, this study reports the first known example of an antisense lncRNA interacting with the product of its sense strand. This binding interaction raises significant scope for newer interaction models possible between head-to-head gene pairs.

## Acknowledgments

The authors acknowledge the contributions of Sourav Ganguly, Priyanshu Das at IIT Gandhinagar, in data collection during a preliminary stage of this work.

## Supplementary

### Competition Assay

The competition assay is performed to determine the role of each G tract present within the G4 forming RNA sequence. Competitor oligonucleotides were designed against each G-tract (Fig. S1). The PQS RNA was incubated with increasing concentrations of competitor oligonucleotide and the sample was run on 15% native PAGE and stained with ThT to visualize the impact of the competitor oligonucleotide on G4 formation (Fig. S2).

**Table S1.**
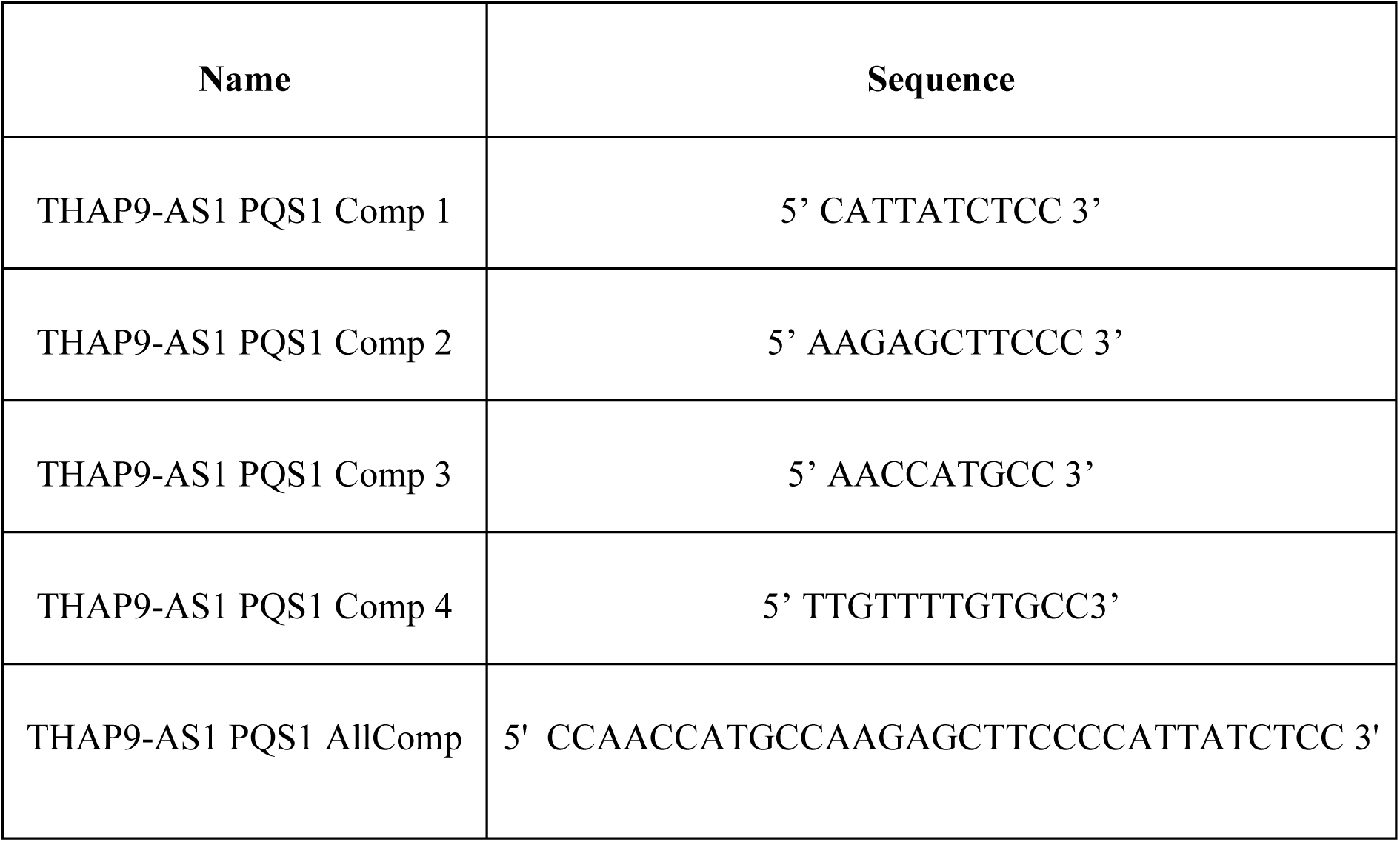
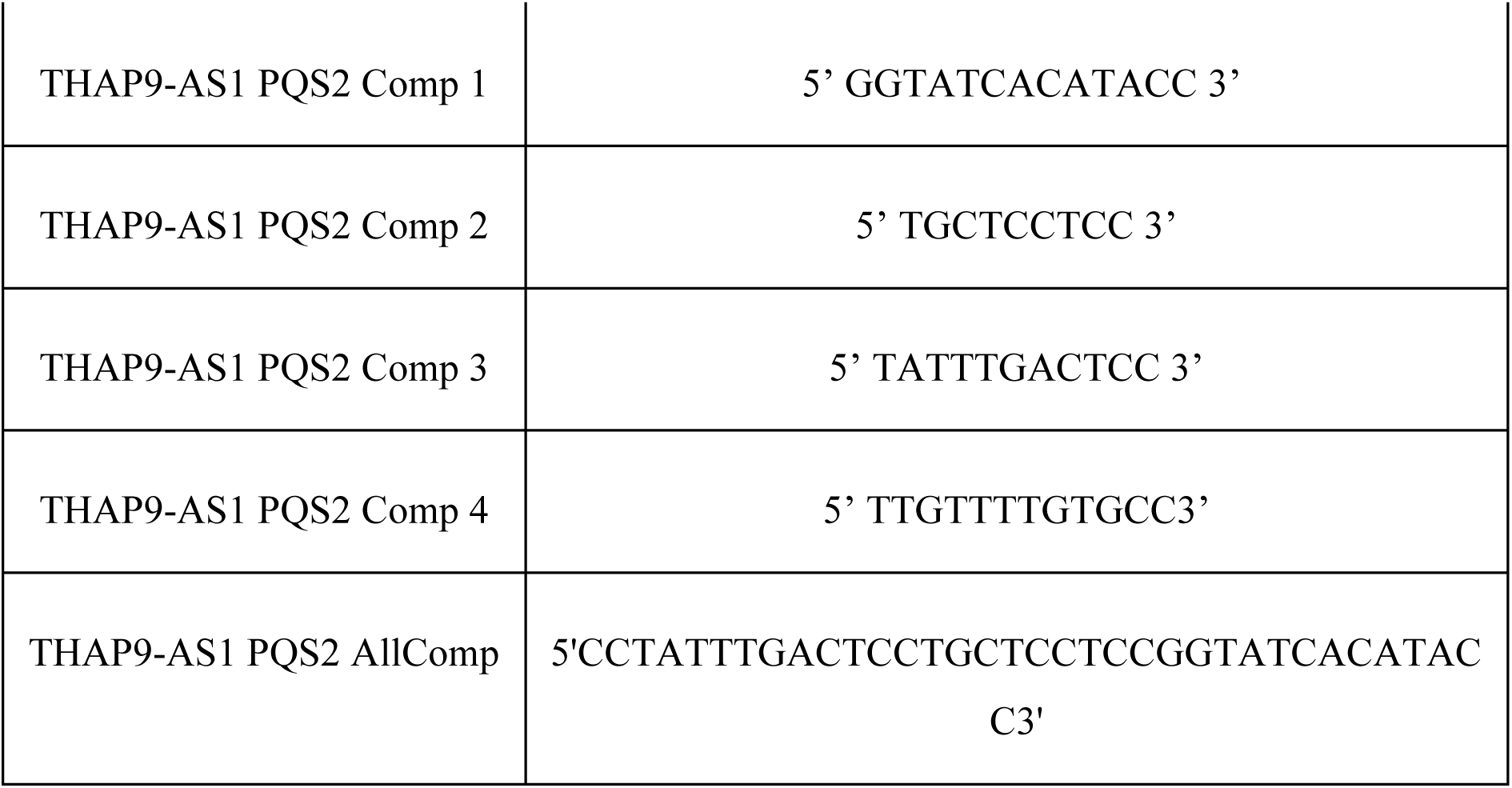
Sequence of competitor oligonucleotides designed for competition assay.

**Figure S1.**
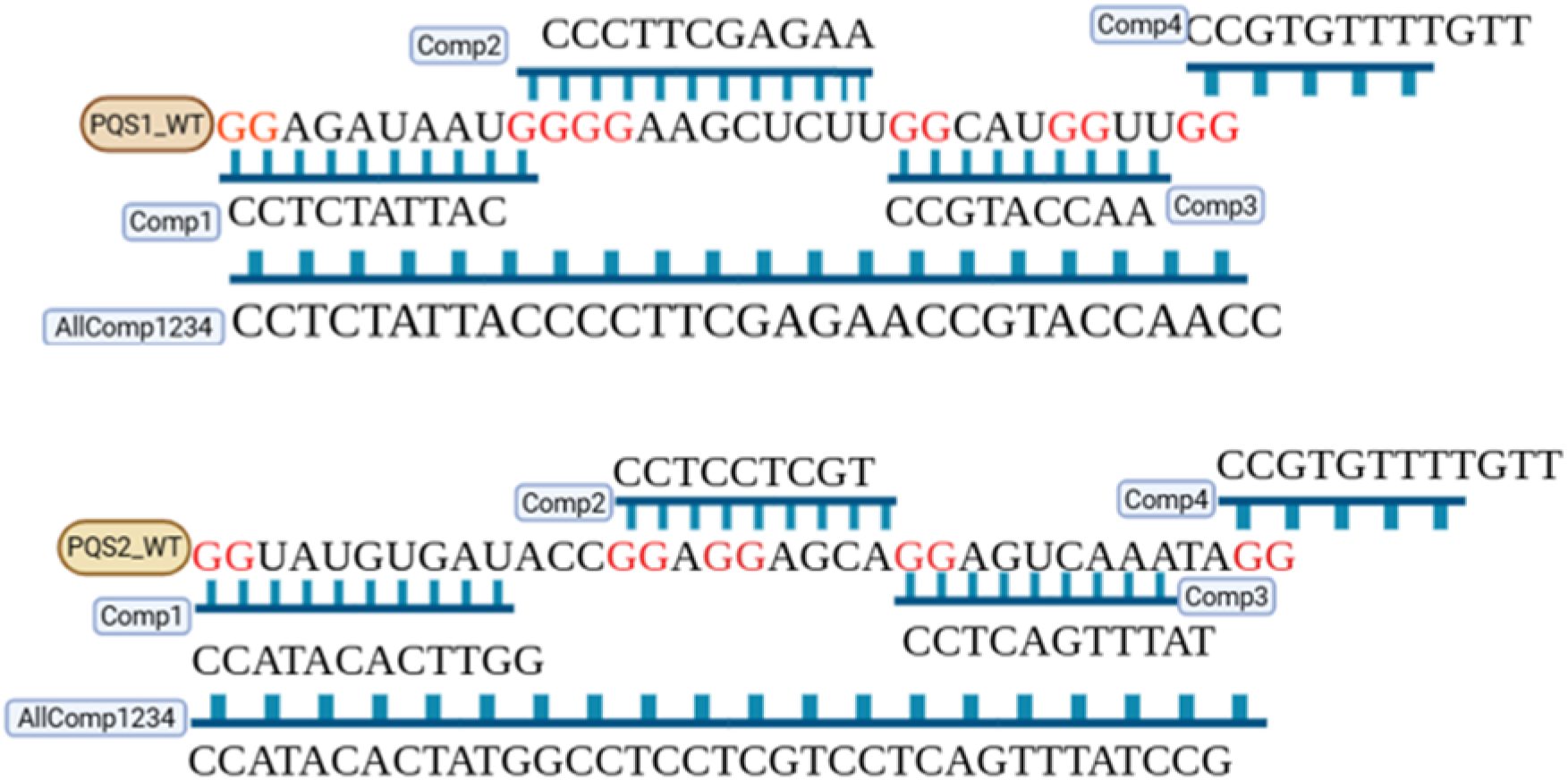
Schematic representation of Competitor oligonucleotides interacting with the G tracts in PQS1 and PQS2

**Figure S2.**
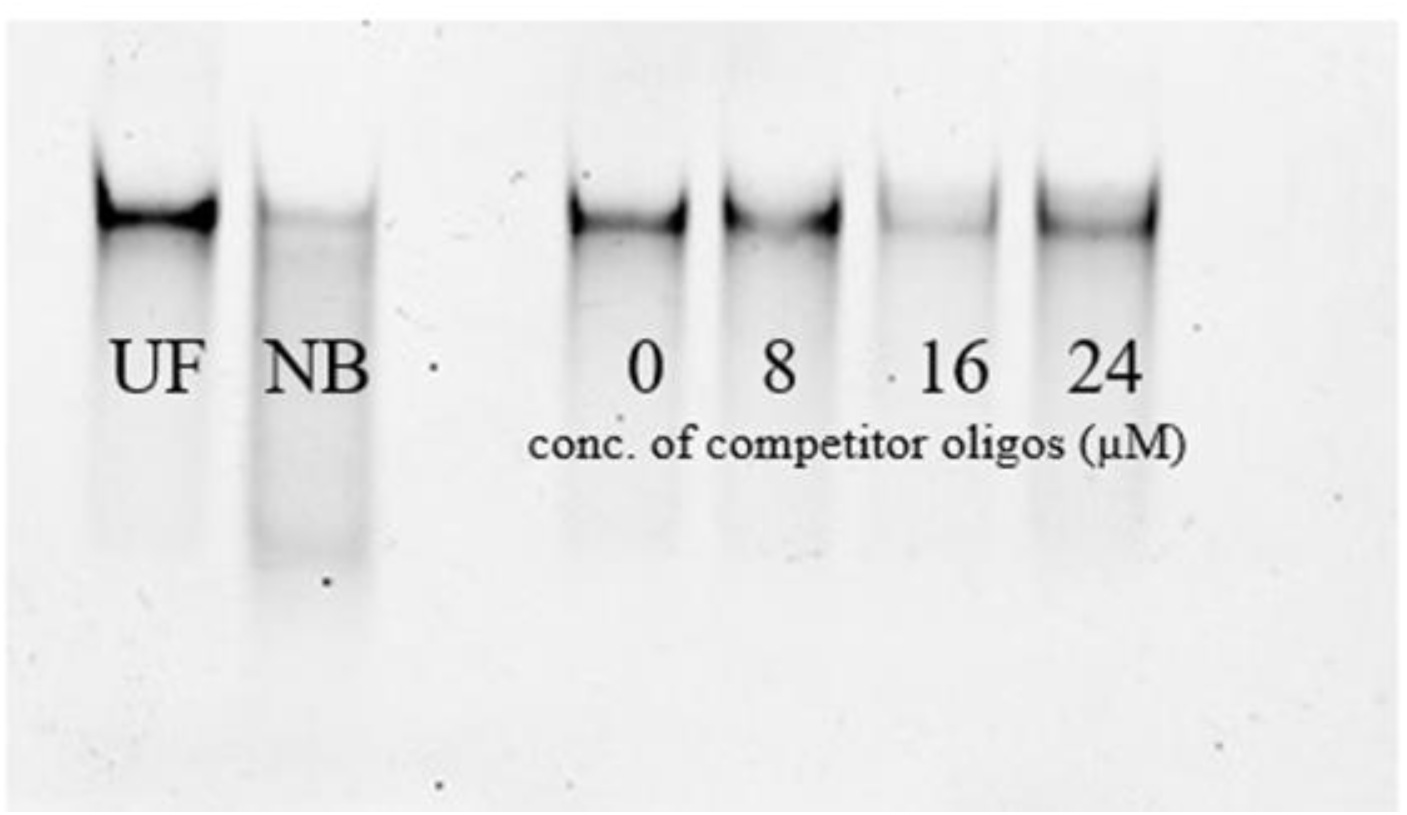
Representative figure showing ThT stained Native gel of competition assay (UF: unfolded RNA; NB: RNA in absence of folding buffer)

**Figure S3.**
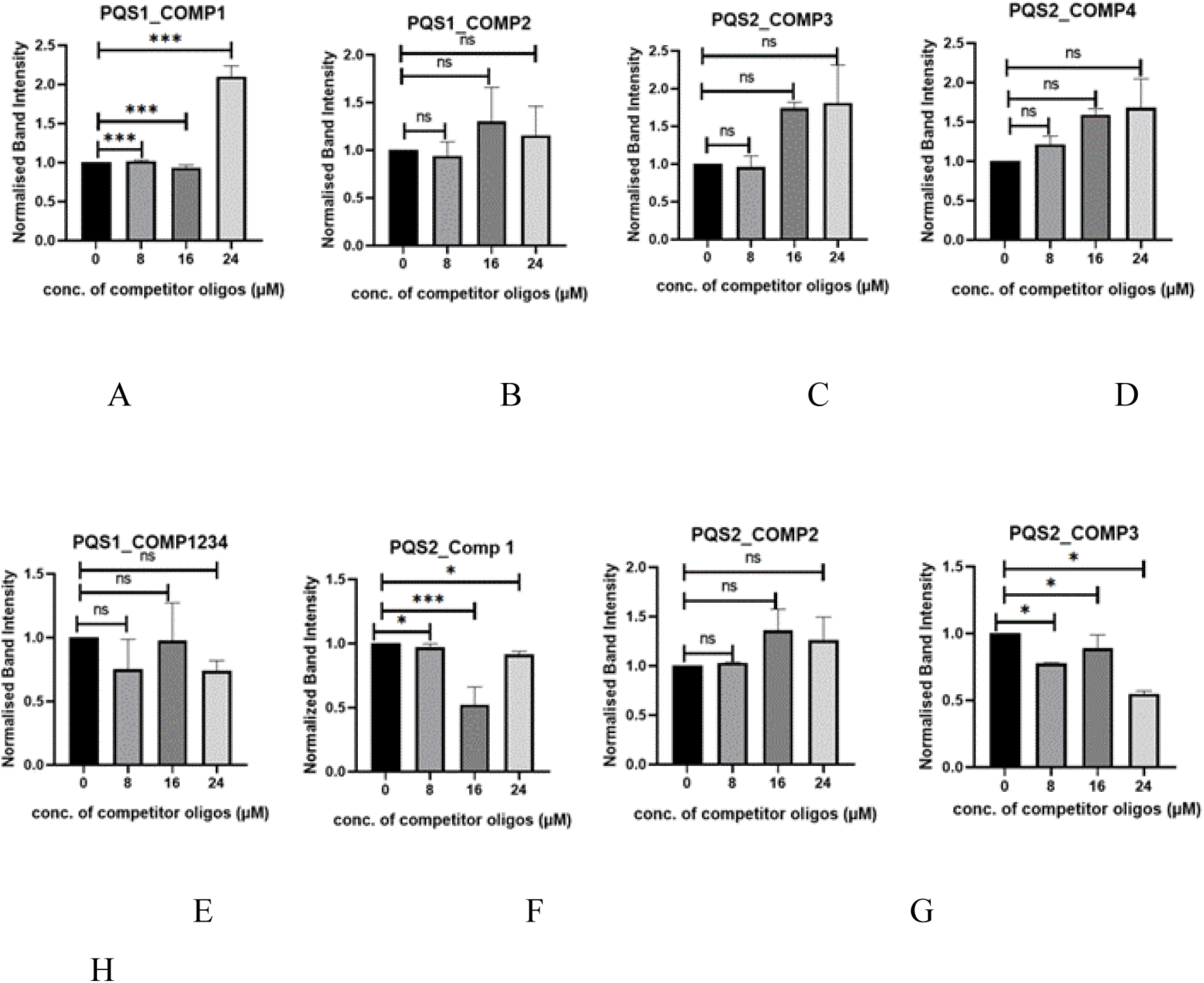
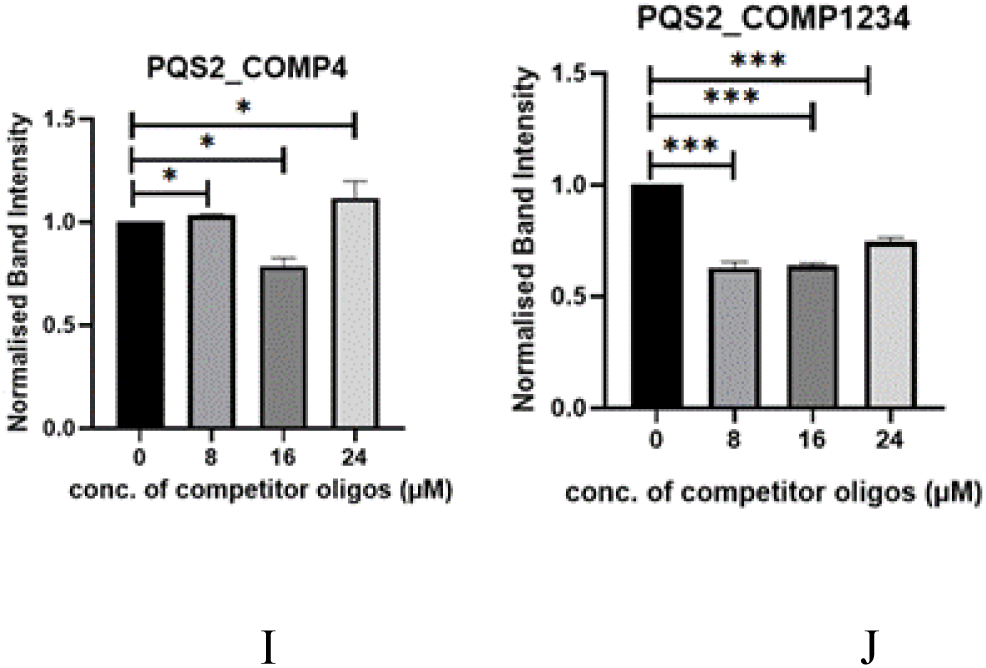
Competition Assay of PQS1 incubated with (A) competitor oligo 1 (B) competitor oligo 2 (C) competitor oligo 3 (D) competitor oligo 4 (E) competitor oligo 1234 (against the whole stretch of PQS). PQS2 incubated with (F) competitor oligo 1 (G) competitor oligo 2 (H) competitor oligo 3 (I) competitor oligo 4 (J) competitor oligo 1234 (against whole stretch of PQS)

Fig. S3 demonstrates that competitor DNA oligos synthesized against individual G tracts within the PQS sequence of PQS1, do not appear to have significant disruptive effects on G4 formation. This could be attributed to the formation of DNA-RNA hybrid G4 complexes.

Even for PQS2, there is marked disruption of G4 formation (Fig. S3 (J)) only in the presence of a competitor oligo against the complete PQS.

**Table S2.**
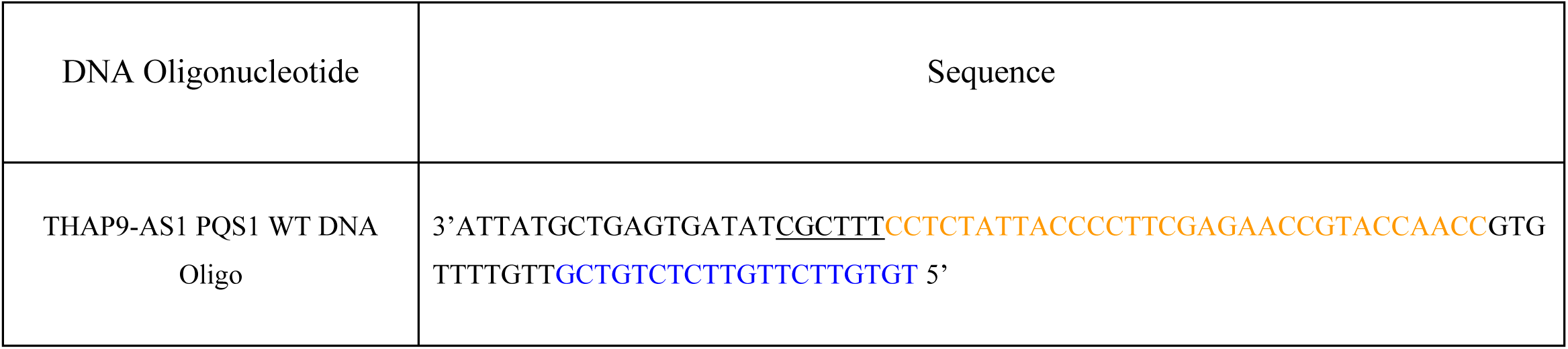
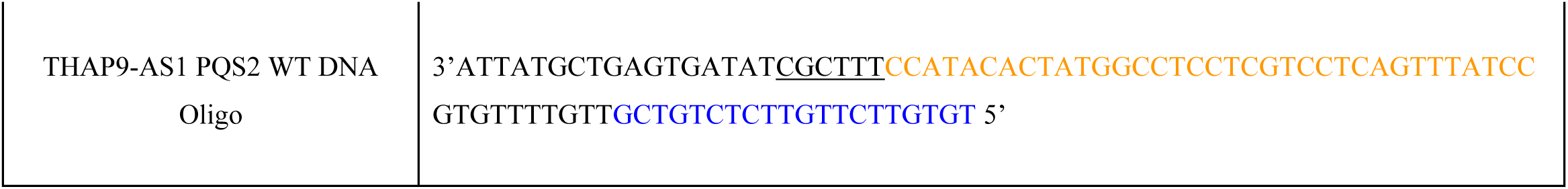
Designed DNA oligonucleotides for IVT of PQS1 and PQS2.

The underlined sequence in each PQS (orange font) represents the extra bases (Red box in Fig. S4) before the PQS and random primer binding site is marked in blue font.

Fig. S4 describes the rationale behind the design.

**Figure S4.**
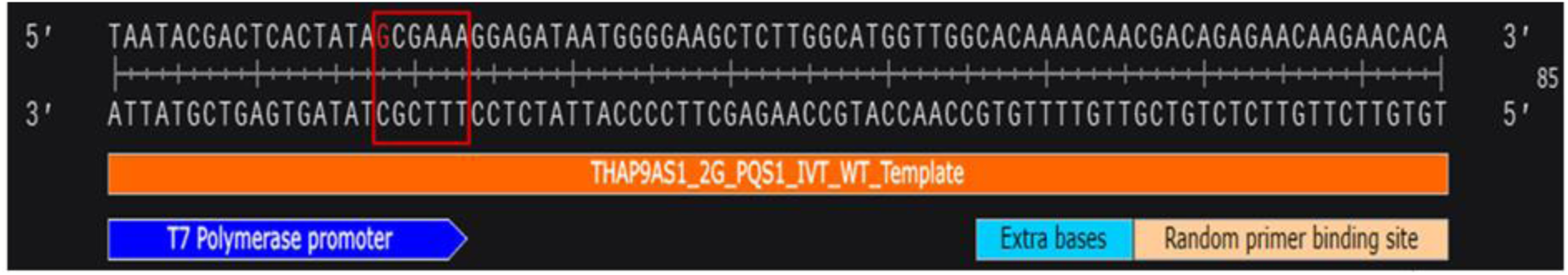
Representative image for PQS1 (GGAGAUAAUGGGGAAGCUCUUGGCAUGGUUGG)

The 3’ to 5’ sequence is the THAP9-AS1_PQS1_WT template DNA oligonucleotide (orange box, synthesised by Sigma) and the 5’ to 3’ sequence corresponds to the actual RNA sequence (with Ts replaced by Us) which will be synthesised in the IVT reaction. The PQS1 is preceded by a T7 polymerase binding site (blue box) as well as extra bases (red box) after the G (highlighted in red font) from where transcription starts so as to prevent addition of the initial G to the rG4 forming sequence (Singh et al., 2023) during IVT. A random primer binding site (peach colour; 5’-TGTGTTCTTGTTCTCTGTCG-3’) is added to facilitate the binding of random primers in subsequent RT stop assays: this site is separated from the PQS by extra bases (cyan) to ensure that when a random primer binds to the RNA, it does not interfere with rG4 formation.

**Table S3.**
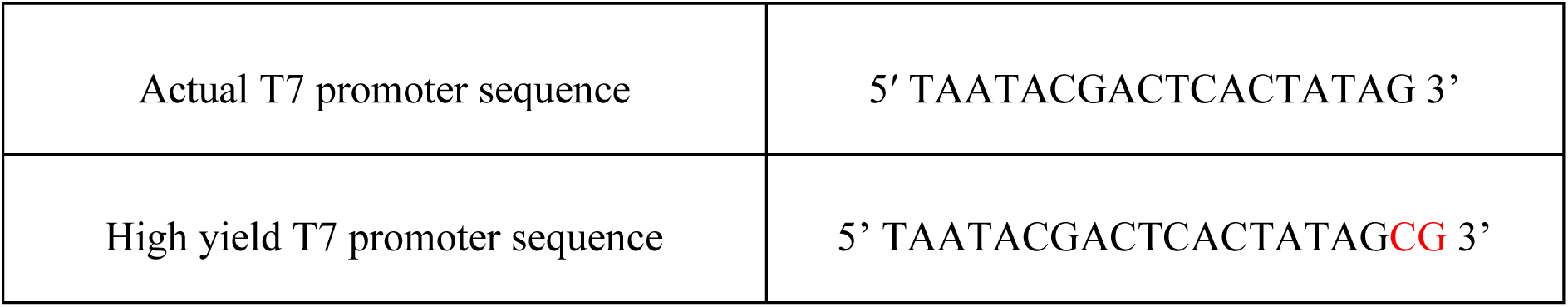
T7 promoter sequence and its modifications.

**Figure S5.**
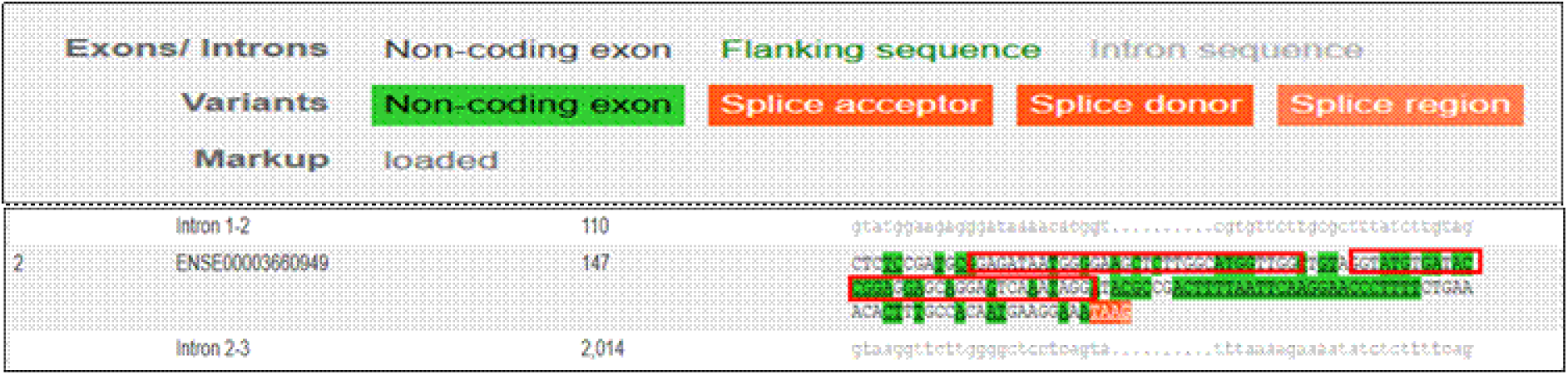
Screenshot from Ensembl showing the presence of PQS1 and PQS2 (marked by red box) separated by 5 nucleotides within the second exon of THAP9-AS1 gene (Transcript ID:<u>ENST00000504520.6</u>)(https://www.ensembl.org/Homo_sapiens/Transcript/Exons?db=core;g=ENSG00000251022;r=4:82893009-82900960;t=ENST00000504520)

**Table S4.**
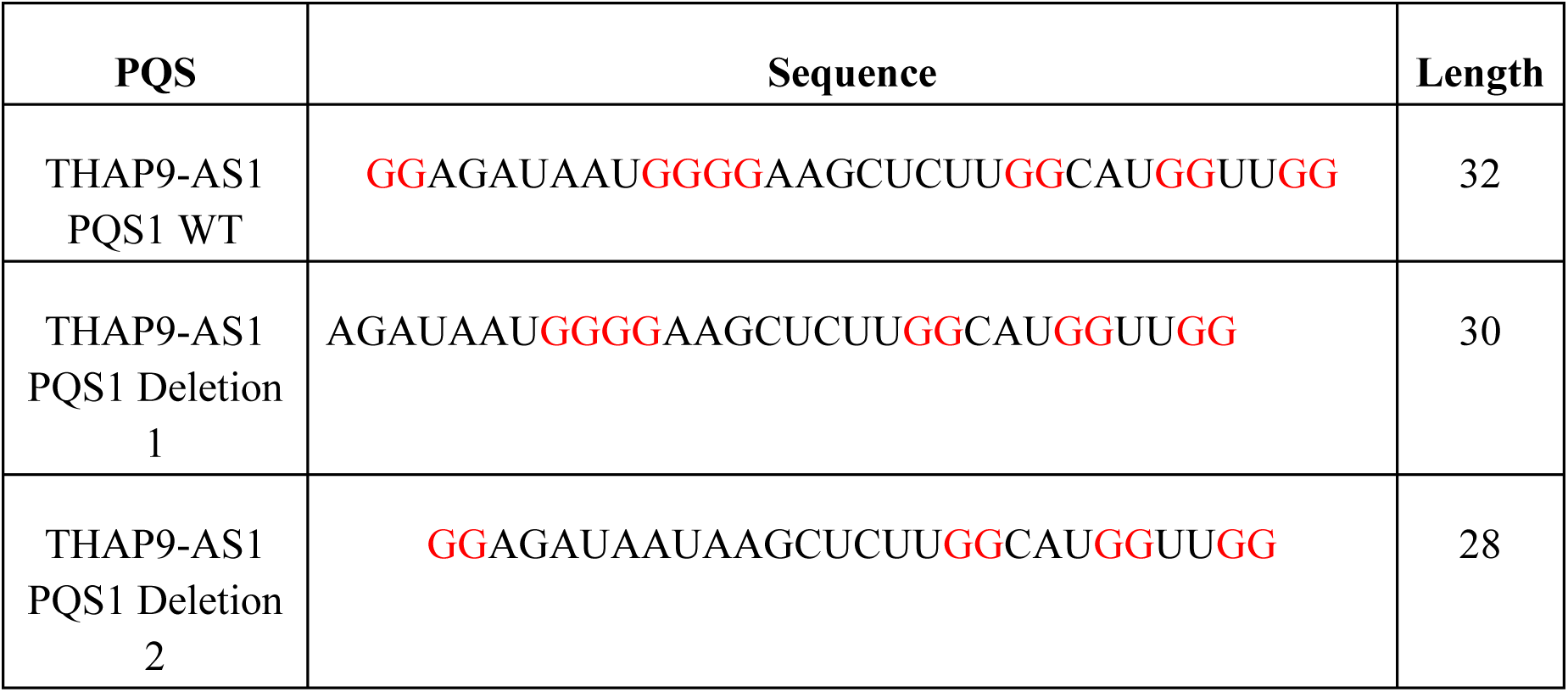
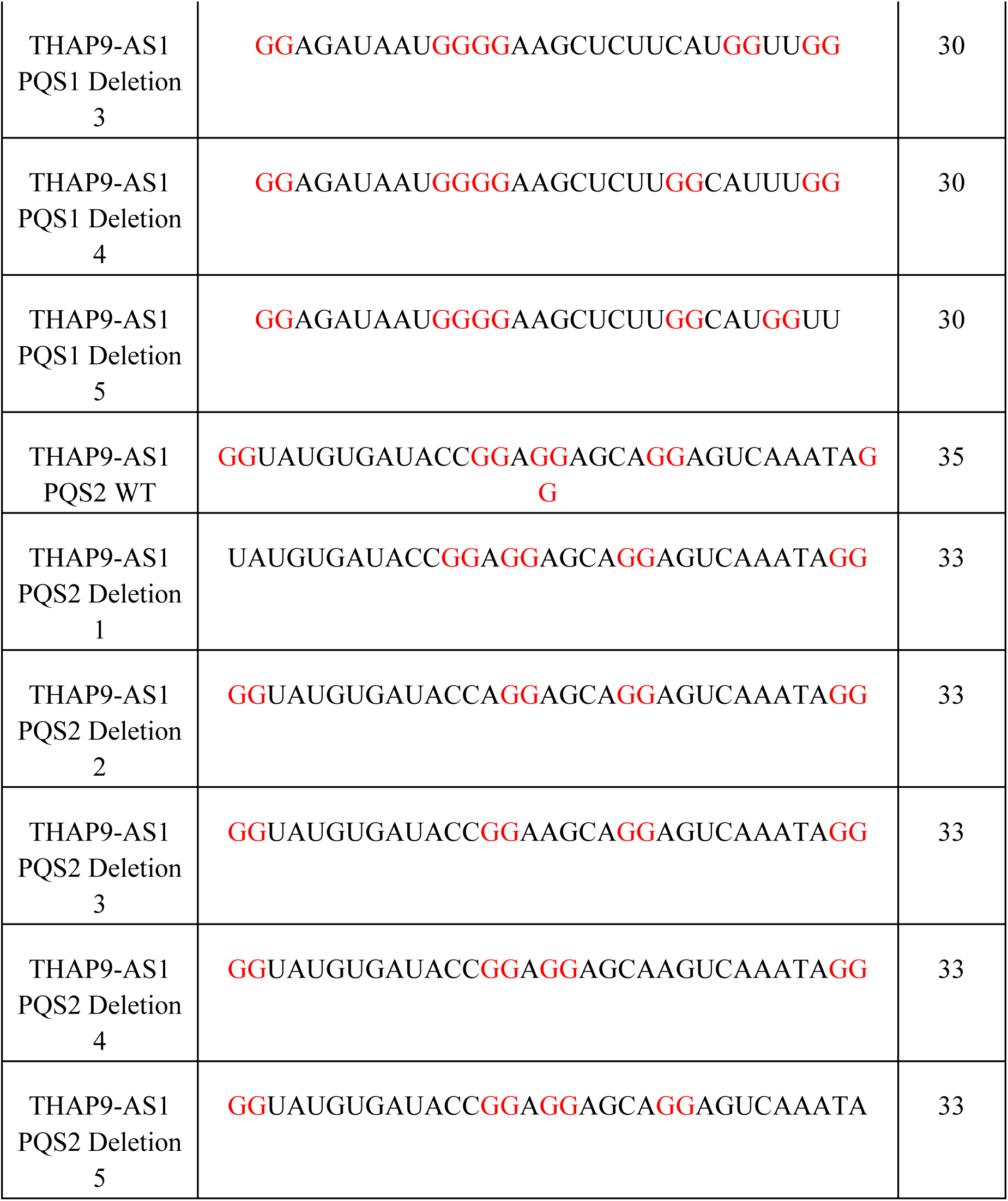
PQS1 and PQS2 WT and deletion mutants synthesized for in vitro studies.

## Bibliography

Albrecht, A. S., & Ørom, U. A. (2016). Bidirectional expression of long ncRNA/protein-coding gene pairs in cancer. Briefings in Functional Genomics, 15, 167–173. 10.1093/bfgp/elv048

Bhat, S. A., Ahmad, S. M., Mumtaz, P. T., Malik, A. A., Dar, M. A., Urwat, U., Shah, R. A., & Ganai, N. A. (2016). Long non-coding RNAs: Mechanism of action and functional utility. Non-Coding RNA Research, 1, 43–50. 10.1016/j.ncrna.2016.11.002

Booy, E. P., Meier, M., Okun, N., Novakowski, S. K., Xiong, S., Stetefeld, J., & McKenna, S. A. (2012). The RNA helicase RHAU (DHX36) unwinds a G4-quadruplex in human telomerase RNA and promotes the formation of the P1 helix template boundary. Nucleic Acids Research, 40(9), 4110–4124. 10.1093/NAR/GKR1306,

Brázda, V., Kolomazník, J., Lýsek, J., Bartas, M., Fojta, M., Šťastný, J., & Mergny, J. L. (2019). G4Hunter web application: a web server for G-quadruplex prediction. Bioinformatics, 35(18), 3493–3495. 10.1093/BIOINFORMATICS/BTZ087

Brown, C. J., Hendrich, B. D., Rupert, J. L., Lafrenière, R. G., Xing, Y., Lawrence, J., & Willard, H. F. (1992). The human XIST gene: Analysis of a 17 kb inactive X-specific RNA that contains conserved repeats and is highly localized within the nucleus. Cell, 71, 527–542. 10.1016/0092-8674(92)90520-M

Cabili, M., Trapnell, C., Goff, L., Koziol, M., Tazon-Vega, B., Regev, A., & Rinn, J. L. (2011). Integrative annotation of human large intergenic noncoding RNAs reveals global properties and specific subclasses. Genes and Development, 25, 1915–1927. 10.1101/gad.17446611

Cheng, J., Ma, H., Yan, M., & Xing, W. (2021). THAP9-AS1/miR-133b/SOX4 positive feedback loop facilitates the progression of esophageal squamous cell carcinoma. Cell Death and Disease, 12(4). 10.1038/s41419-021-03690-z

Cléry, A., Allain, F. H.-T., & Antoine Cléry, F. H.-T. A. (2013). FROM STRUCTURE TO FUNCTION OF RNA BINDING DOMAINS. In https://www.ncbi.nlm.nih.gov/books/NBK63528/.

De La Faverie, A. R., Guédin, A., Bedrat, A., Yatsunyk, L. A., & Mergny, J. L. (2014). Thioflavin T as a fluorescence light-up probe for G4 formation. Nucleic Acids Research, 42(8), e65–e65. 10.1093/NAR/GKU111

del Villar-Guerra, R., Trent, J. O., & Chaires, J. B. (2018). G-Quadruplex Secondary Structure Obtained from Circular Dichroism Spectroscopy. Angewandte Chemie - International Edition, 57, 7171–7175. 10.1002/anie.201709184

Fang, Y., & Fullwood, M. J. (2016). Roles, Functions, and Mechanisms of Long Non-coding RNAs in Cancer. In Genomics, Proteomics and Bioinformatics (Vol. 14, pp. 42–54). Beijing Genomics Institute. 10.1016/j.gpb.2015.09.006

Fang, Y., Liu, X., Liu, Y., & Xu, N. (2024). Insights into the Mode and Mechanism of Interactions Between RNA and RNA-Binding Proteins. In International Journal of Molecular Sciences (Vol. 25). Multidisciplinary Digital Publishing Institute (MDPI). 10.3390/ijms252111337

Fernandes, J. C. R., Acuña, S. M., Aoki, J. I., Floeter-Winter, L. M., & Muxel, S. M. (2019). Long non-coding RNAs in the regulation of gene expression: Physiology and disease. In Non-coding RNA (Vol. 5). MDPI AG. 10.3390/ncrna5010017

Gerstberger, S., Hafner, M., & Tuschl, T. (2014). A census of human RNA-binding proteins. Nature Reviews Genetics, 15, 829–845. 10.1038/nrg3813

Grammatikakis, I., & Lal, A. (2021). Significance of lncRNA abundance to function. Mammalian Genome 2021 33:2, 33(2), 271–280. 10.1007/S00335-021-09901-4

Haase, A. D., Ketting, R. F., Lai, E. C., van Rij, R. P., Siomi, M., Svoboda, P., van Wolfswinkel, J. C., & Wu, P. H. (2024). PIWI-interacting RNAs: who, what, when, where, why, and how. In EMBO Journal (Vol. 43, pp. 5335–5339). EMBO Press. 10.1038/s44318-024-00253-8

Hao, A., Wang, Y., Stovall, D. B., Wang, Y., & Sui, G. (2021). Emerging Roles of LncRNAs in the EZH2-regulated Oncogenic Network. International Journal of Biological Sciences, 17(13), 3268. 10.7150/IJBS.63488

Hermann, T., & Patel, D. J. (2000). RNA bulges as architectural and recognition motifs. In Structure (Vol. 8). Current Biology Ltd. 10.1016/S0969-2126(00)00110-6

Hou, J., Zhang, G., Wang, X., Wang, Y., & Wang, K. (2023). Functions and mechanisms of lncRNA MALAT1 in cancer chemotherapy resistance. Biomarker Research, 11, 23. 10.1186/s40364-023-00467-8

Jia, W., Zhang, J., Ma, F., Hao, S., Li, X., Guo, R., Gao, Q., Sun, Y., Jia, J., & Li, W. (2019). Long noncoding RNA THAP9-AS1 is induced by Helicobacter pylori and promotes cell growth and migration of gastric cancer. OncoTargets and Therapy, 12, 6653. 10.2147/OTT.S201832

Katayama, S., Tomaru, Y., Kasukawa, T., Waki, K., Nakanishi, M., Nakamura, M., Nishida, H., Yap, C. C., Suzuki, M., Kawai, J., Suzuki, H., Carninci, P., Hayashizaki, Y., Wells, C., Frith, M., Ravasi, T., Pang, K. C., Hallinan, J., Mattick, J., … Wahlestedt, C. (2005). Molecular biology: Antisense transcription in the mammalian transcriptome. Science, 309, 1564–1566. 10.1126/science.1112009

Kikin, O., D’Antonio, L., & Bagga, P. S. (2006). QGRS Mapper: a web-based server for predicting G-quadruplexes in nucleotide sequences. Nucleic Acids Research, 34(suppl_2), W676–W682. 10.1093/NAR/GKL253

Kim, J., Cheong, C., & Moore, P. B. (1991). Tetramerization of an RNA oligonucleotide containing a GGGG sequence. Nature, 351, 331–332. 10.1038/351331a0

Kim, T. K., Hemberg, M., Gray, J. M., Costa, A. M., Bear, D. M., Wu, J., Harmin, D. A., Laptewicz, M., Barbara-Haley, K., Kuersten, S., Markenscoff-Papadimitriou, E., Kuhl, D., Bito, H., Worley, P. F., Kreiman, G., & Greenberg, M. E. (2010). Widespread transcription at neuronal activity-regulated enhancers. Nature, 465, 182–187. 10.1038/nature09033

Lee, S., Kopp, F., Chang, T. C., Sataluri, A., Chen, B., Sivakumar, S., Yu, H., Xie, Y., & Mendell, J. T. (2016). Noncoding RNA NORAD Regulates Genomic Stability by Sequestering PUMILIO Proteins. Cell, 164(1–2), 69–80. 10.1016/j.cell.2015.12.017

Li, N., Yang, G., Luo, L., Ling, L., Wang, X., Shi, L., Lan, J., Jia, X., Zhang, Q., Long, Z., Liu, J., Hu, W., He, Z., Liu, H., Liu, W., & Zheng, G. (2020). lncRNA THAP9-AS1 Promotes Pancreatic Ductal Adenocarcinoma Growth and Leads to a Poor Clinical Outcome via Sponging miR-484 and Interacting with YAP. Clinical Cancer Research, 26(7), 1736–1748. 10.1158/1078-0432.CCR-19-0674

Mattick, J. S., Amaral, P. P., Carninci, P., Carpenter, S., Chang, H. Y., Chen, L. L., Chen, R., Dean, C., Dinger, M. E., Fitzgerald, K. A., Gingeras, T. R., Guttman, M., Hirose, T., Huarte, M., Johnson, R., Kanduri, C., Kapranov, P., Lawrence, J. B., Lee, J. T., … Wu, M. (2023a). Long non-coding RNAs: definitions, functions, challenges and recommendations. Nature Reviews Molecular Cell Biology, 24(6), 430–447. 10.1038/s41580-022-00566-8

Mattick, J. S., Amaral, P. P., Carninci, P., Carpenter, S., Chang, H. Y., Chen, L. L., Chen, R., Dean, C., Dinger, M. E., Fitzgerald, K. A., Gingeras, T. R., Guttman, M., Hirose, T., Huarte, M., Johnson, R., Kanduri, C., Kapranov, P., Lawrence, J. B., Lee, J. T., … Wu, M. (2023b). Long non-coding RNAs: definitions, functions, challenges and recommendations. Nature Reviews Molecular Cell Biology, 24, 430–447. 10.1038/s41580-022-00566-8

Mei, Y., Deng, Z., Vladimirova, O., Gulve, N., Johnson, F. B., Drosopoulos, W. C., Schildkraut, C. L., & Lieberman, P. M. (2021). TERRA G-quadruplex RNA interaction with TRF2 GAR domain is required for telomere integrity. Scientific Reports, 11. 10.1038/s41598-021-82406-x

Meier-Stephenson, V. (2022). G4-quadruplex-binding proteins: review and insights into selectivity. Biophysical Reviews, 14, 635–654. 10.1007/s12551-022-00952-8

Meller, V. H., & Rattner, B. P. (2002). The roX genes encode redundant male-specific lethal transcripts required for targeting of the MSL complex. EMBO Journal, 21, 1084–1091. 10.1093/emboj/21.5.1084

Mohanty, J., Barooah, N., Dhamodharan, V., Harikrishna, S., Pradeepkumar, P. I., & Bhasikuttan, A. C. (2013). Thioflavin T as an efficient inducer and selective fluorescent sensor for the human telomeric G-quadruplex DNA. Journal of the American Chemical Society, 135(1), 367–376. 10.1021/JA309588H

Muppirala, U. K., Honavar, V. G., & Dobbs, D. (2011). Predicting RNA-Protein Interactions Using Only Sequence Information. BMC Bioinformatics 2011 12:1, 12(1), 489-. 10.1186/1471-2105-12-489

Obuse, C., & Hirose, T. (2023). Functional domains of nuclear long noncoding RNAs: Insights into gene regulation and intracellular architecture. In Current Opinion in Cell Biology (Vol. 85). Elsevier Ltd. 10.1016/j.ceb.2023.102250

Pachnis, V., Belayew, A., & Tilghman, S. M. (1984). Locus unlinked to α-fetoprotein under the control of the murine raf and Rif genes. Proceedings of the National Academy of Sciences of the United States of America, 81(17 I), 5523–5527. 10.1073/pnas.81.17.5523

Pelechano, V., & Steinmetz, L. M. (2013). Gene regulation by antisense transcription. Nature Reviews Genetics, 14(12), 880–893. 10.1038/nrg3594

Qu, Z., & Adelson, D. L. (2012). Evolutionary conservation and functional roles of ncRNA. Frontiers in Genetics, 3(OCT), 35289. 10.3389/FGENE.2012.00205/TEXT

Rashmi, R., & Majumdar, S. (2022). Pan-Cancer Analysis Reveals the Prognostic Potential of the THAP9/THAP9-AS1 Sense–Antisense Gene Pair in Human Cancers. Non-Coding RNA, 8(4). 10.3390/ncrna8040051

Rinn, J. L., & Chang, H. Y. (2012). Genome regulation by long noncoding RNAs. Annual Review of Biochemistry, 81, 145–166. 10.1146/annurev-biochem-051410-092902

Ryder, S. P., Recht, M. I., & Williamson, J. R. (2008). Quantitative analysis of protein-RNA interactions by gel mobility shift. Methods in Molecular Biology (Clifton, N.J.), 488, 99–115. 10.1007/978-1-60327-475-3_7

Salmena, L., Poliseno, L., Tay, Y., Kats, L., & Pandolfi, P. P. (2011). A ceRNA hypothesis: The rosetta stone of a hidden RNA language? In Cell (Vol. 146, pp. 353–358). Elsevier B.V. 10.1016/j.cell.2011.07.014

Santos, T., Salgado, G. F., Cabrita, E. J., & Cruz, C. (2021). G-Quadruplexes and Their Ligands: Biophysical Methods to Unravel G-Quadruplex/Ligand Interactions. Pharmaceuticals 2021, Vol. 14, Page 769, 14(8), 769. 10.3390/PH14080769

Simko, E. A. J., Liu, H., Zhang, T., Velasquez, A., Teli, S., Haeusler, A. R., & Wang, J. (2020). G-quadruplexes offer a conserved structural motif for NONO recruitment to NEAT1 architectural lncRNA. Nucleic Acids Research, 48(13), 7421–7438. 10.1093/NAR/GKAA475

Singh, D., Desai, N., Shah, V., & Datta, B. (2023). In Silico Identification of Potential Quadruplex Forming Sequences in LncRNAs of Cervical Cancer. International Journal of Molecular Sciences, 24. 10.3390/ijms241612658

Statello, L., Guo, C. J., Chen, L. L., & Huarte, M. (2021). Gene regulation by long non-coding RNAs and its biological functions. In Nature Reviews Molecular Cell Biology (Vol. 22, pp. 96–118). Nature Research. 10.1038/s41580-020-00315-9

Su, Y., Xie, R., & Xu, Q. (2022a). LncRNA THAP9-AS1 highly expressed in tissues of hepatocellular carcinoma and accelerates tumor cell proliferation. Clinics and Research in Hepatology and Gastroenterology, 46(10), 102025. 10.1016/J.CLINRE.2022.102025

Su, Y., Xie, R., & Xu, Q. (2022b). LncRNA THAP9-AS1 highly expressed in tissues of hepatocellular carcinoma and accelerates tumor cell proliferation. Clinics and Research in Hepatology and Gastroenterology, 46. 10.1016/j.clinre.2022.102025

Varizhuk, A., Ischenko, D., Smirnov, I., Tatarinova, O., Severov, V., Novikov, R., Tsvetkov, V., Naumov, V., Kaluzhny, D., & Pozmogova, G. (2014). An Improved Search Algorithm to Find G-Quadruplexes in Genome Sequences. BioRxiv. 10.1101/001990

Villegas, V. E., & Zaphiropoulos, P. G. (2015). Neighboring Gene Regulation by Antisense Long Non-Coding RNAs. International Journal of Molecular Sciences, 16(2), 3251. 10.3390/IJMS16023251

Vorlíčková, M., Bednářová, K., Kejnovská, I., & Kypr, J. (2007). Intramolecular and intermolecular guanine quadruplexes of DNA in aqueous salt and ethanol solutions. Biopolymers, 86, 1–10. 10.1002/bip.20672

Woolard, E., & Chorley, B. N. (2018). The role of noncoding RNAs in gene regulation. In Toxicoepigenetics: Core Principles and Applications (pp. 217–235). Elsevier. 10.1016/B978-0-12-812433-8.00009-5

Wu, Y., Yin, Q., Zhang, X., Zhu, P., Luan, H., & Chen, Y. (2020). Long Noncoding RNA THAP9-AS1 and TSPOAP1-AS1 Provide Potential Diagnostic Signatures for Pediatric Septic Shock. BioMed Research International, 2020, 7170464. 10.1155/2020/7170464

Wu, Y., Zhang, L., Zhang, L., Wang, Y., Li, H., Ren, X., Wei, F., Yu, W. W., Liu, T., Wang, X., Zhou, X., Yu, J., & Hao, X. (2015). Long non-coding RNA HOTAIR promotes tumor cell invasion and metastasis by recruiting EZH2 and repressing E-cadherin in oral squamous cell carcinoma. International Journal of Oncology, 46, 2586–2594. 10.3892/ijo.2015.2976

Xu, S., Li, Q., Xiang, J., Yang, Q., Sun, H., Guan, A., Wang, L., Liu, Y., Yu, L., Shi, Y., Chen, H., & Tang, Y. (2016). Thioflavin T as an efficient fluorescence sensor for selective recognition of RNA G-quadruplexes. Scientific Reports, 6. 10.1038/srep24793

Yamazaki, T., Souquere, S., Chujo, T., Kobelke, S., Chong, Y. S., Fox, A. H., Bond, C. S., Nakagawa, S., Pierron, G., & Hirose, T. (2018). Functional Domains of NEAT1 Architectural lncRNA Induce Paraspeckle Assembly through Phase Separation. Molecular Cell, 70(6), 1038–1053.e7. 10.1016/j.molcel.2018.05.019

Zaccaria, F., & Fonseca Guerra, C. (2018). RNA versus DNA G-Quadruplex: The Origin of Increased Stability. Chemistry - A European Journal, 24, 16315–16322. 10.1002/chem.201803530

Zhang, J., Chen, B., Gan, C., Sun, H., Zhang, J., & Feng, L. (2023). A Comprehensive Review of Small Interfering RNAs (siRNAs): Mechanism, Therapeutic Targets, and Delivery Strategies for Cancer Therapy. In International Journal of Nanomedicine (Vol. 18, pp. 7605–7635). Dove Medical Press Ltd. 10.2147/IJN.S436038

Zhao, J., Sun, B. K., Erwin, J. A., Song, J. J., & Lee, J. T. (2008). Polycomb proteins targeted by a short repeat RNA to the mouse X chromosome. Science, 322, 750–756. 10.1126/science.1163045

